# Translational Opportunity of Engineered IFNγ-eEVs Through Targeted Inhibition of JAK/STAT1 Signaling, Mimicking IVIg Therapy

**DOI:** 10.64898/2026.04.29.721601

**Authors:** Kylie E. Preihs, Kubra Karagoz, Claire R. Shuey, Amritha Achuthkumar, Ana M. Pivovarnik, Sophia M. Crocker, Michelle L. Pleet, Jubin George, Robert D. Carlson, Adam E. Snook, Adam J. Luginbuhl, Peter J. Wermuth, Andreas Möller, Jennifer C. Jones, Larry A. Harshyne, Alice P. Pentland, Mỹ G. Mahoney

## Abstract

Immunoglobulin (Ig) replacement therapies (IgRT) including intravenous (IVIg) and subcutaneous (SCIg), are pooled IgG preparations widely used to restore humoral immunity and to suppress pathological inflammation in autoimmune and inflammatory disorders. Despite broad clinical use, the mechanisms underlying their immunomodulatory effects remain incompletely defined. Here, we identify extracellular vesicle (EV)-associated cytokines as mediators of IVIg activity. Multiplex bead-based flow cytometry revealed that EVs isolated by size exclusion followed by ultracentrifugation from IVIg were CD63 positive but depleted of platelet-derived and HLA markers relative to EVs from unprocessed human plasma. Luminex profiling demonstrated substantial reduction of pro-inflammatory cytokines in IVIg EVs. Notably, although IVIg EVs contained abundant IFNγ, they failed to activate IFNGR/JAK/STAT1 signaling. Instead, prolonged exposure to IVIg EVs suppressed subsequent IFNγ-induced STAT1 activation. Engineered IFNγ-coated EVs (IFNγ-eEVs) recapitulated both activating and inhibitory effects indicating context-dependent signaling bias. Critically, cold ethanol precipitation, a key step in IVIg manufacturing, selectively abrogated the activating function of IFNγ-eEVs while preserving their inhibitory capacity. These findings define a previously unrecognized mechanism where IVIg processing generates EVs that bias IFNγ signaling toward suppression. EV-associated cytokines therefore represent a generalizable pathway through which IVIg exerts anti-inflammatory effects across immune-mediated diseases.

## 1 Introduction

Immunoglobulin (Ig) replacement therapy (IgRT) marked a pivotal advancement in the management of immune deficiencies, reflecting decades of therapeutic innovation and refinement (Ahmed et al., 2025; Späth et al., 2017; von Gunten et al., 2023). The IgRT field has progressed substantially since the introduction of intramuscular Ig preparations in the 1950s, with the subsequent approval of intravenous immunoglobulin (IVIg) gaining FDA approval in 1981 and subcutaneous immunoglobulin (SCIg) in 2006. These advances have expanded treatment accessibility and efficacy, particularly among aging populations with increased susceptibility to immune-related pathologies. Contemporary IVIg preparations consist predominantly of rigorously purified polyclonal antibodies (>95% IgG) isolated from the pooled plasma of up to 15,000 healthy plasma donors per lot, providing a broad and diverse immunological repertoire (Jolles et al., 2005).

Clinically, IVIg is widely used in the treatment of primary and secondary immunodeficiencies, as well as a range of autoimmune and inflammatory diseases (Jolles et al., 2005; Nimmerjahn and Ravetch, 2007; Sewell and Jolles, 2002). Its therapeutic efficacy, including high response rates approaching 75% in conditions such as autoimmune thrombocytopenia (Hsia et al., 2015), is attributed to both immune replacement and immunomodulatory functions. Mechanistically, IVIg exerts diverse and multifaceted effects, by supplying a diverse repertoire of protective antibodies that bolster immunity in immunocompromised patients while neutralizing pathogenic autoantibodies (Bayry et al., 2023). High-dose IVIg also rapidly alleviates acute inflammatory responses (Tjon et al., 2015), potentially through interference with the complement cascade (Basta, 1996), modulation of FCγ receptors, or sialylation of the IgG Fc region (Li et al., 2021). Despite these well-documented effects, the precise molecular and cellular mechanisms underlying IVIg’s activity, particularly in chronic inflammatory states, remain incompletely defined, limiting the development of more targeted and optimized Ig-based interventions with enhanced specificity, improved efficacy, and reduced adverse effects (Schwab and Nimmerjahn, 2013).

Extracellular vesicles (EVs) have emerged as critical mediators of intercellular communication, particularly in the immune system (Welsh et al., 2024b). These membrane-bound nanoparticles, present in all biological fluids including plasma, transport bioactive cargos including lipids, proteins, and nucleic acids (Johnsen et al., 2019; Welsh et al., 2024b). Notably, EVs may display Fc receptors such as CD16, enabling potential IgG binding through protein-protein interactions (Hofmann et al., 2020). EVs influence recipient cells either through direct surface engagement or by delivering internalized cargo, thereby modulating recipient cellular function and immune function (Flemming et al., 2024; Jackson Cullison et al., 2024). EVs serve as promising biomarkers for disease monitoring and as key immune modulators (Buzas, 2023). A notable EV feature is the biomolecular corona (BC), an adsorbed layer of functional biomolecules from plasma and interstitial fluids that can modulate inflammatory signaling (Duong et al., 2019; Hosseini et al., 2022; Liang et al., 2023a; Tóth et al., 2021; Wolf et al., 2022). Cytokines and their receptors have been identified within the EV BC, highlighting a potential role for EVs in cytokine presentation and regulation (Cossetti et al., 2014; Esmaeili et al., 2025; Lima et al., 2021).

Among these cytokines, interferon gamma (IFNγ), a ∼25 kDa pro-inflammatory cytokine secreted primarily by T and NK cells, plays a central role in orchestrating immune activation, driving differentiation and activation of immune cells including B cells and macrophages (Casanova et al., 2024; Ivashkiv, 2018). IFNγ induces production of chemokines like CXCL10 (IP-10), coordinating innate and adaptive immunity (Dufour et al., 2002). IFNγ displayed on EVs can activate STAT1 signaling via binding to IFNγ receptor 1 (IFNGR1) on target cells (Cossetti et al., 2014). However, IFNγ levels must be tightly regulated, as sustained exposure triggers negative feedback pathways leading to T cell exhaustion, immune suppression, and resolution of inflammation (Andrews et al., 2024; Benci et al., 2019; Hosking et al.; Mazet et al., 2023; Michalska et al., 2018). IFNγ thus exhibits context-dependent dual roles in inflammation. Importantly, IVIg has been shown to interfere with IFNγ signaling pathways in macrophages and T cells, potentially through downregulation of IFNGR2, suggesting a possible intersection between Ig-based therapies and cytokine regulation (Park-Min et al., 2007).

Despite the established abundance of EVs in plasma, their presence and functional relevance in processed IVIg products have not been previously characterized. In this study, we investigate the persistence, composition, and immunological activity of EVs isolated from IVIg in comparison to unprocessed human plasma (UHP). Using size-exclusion chromatography (SEC) and/or differential ultracentrifugation (dUC), we demonstrate that IVIg contains intact EV populations with distinct cytokine profiles and immunomodulatory properties. Our findings reveal that IVIg-derived EVs suppress IFNγ-receptor activation, uncovering a novel mechanism underlying IVIg’s anti-inflammatory activity. These results were corroborated using engineered IFNγ-eEVs processed by cold ethanol precipitation (cEP), the initial step in IVIg processing. This represents the first demonstration of intact, functional EVs in therapeutic IVIg preparations capable of suppressing inflammatory pathways by surface-associated cytokine inactivation. These results provide new insight into the complexity of IVIg therapeutics and identify EV-associated pathways as potential targets for next-generation immunomodulatory strategies.

## 2 Materials and Methods

### 2.1 Plasma Procurement

Individual unprocessed human plasma samples (UHPi) from healthy donors were obtained with informed consent from the TJU Biorepository. Briefly, whole blood was collected in tubes treated with the anticoagulant EDTA and centrifuged at 300 x g for 20 min. The buffy coat and peripheral blood cells were discarded while the plasma supernatant was collected and stored at -80°C. Rochester Medical Center, Rochester, NY. Pooled unprocessed human plasma (UHPp) was obtained from Innovative Research (Novi, MI). IVIg (10%; Privigen^®^, King of Prussia, PA) and SCIg (20% liquid; Hizentra^®^, Kankakee, IL) were obtained from the Department of Medicine: Allergy, Immunology, and Rheumatology, University of Rochester Medical Center, Rochester, NY.

### 2.2 Purification and Characterization of Extracellular Vesicles

Plasma EV isolation was performed by SEC as previously described using qEV1 or qEV10 columns (IZON Science, Medford, MA) and 1 or 5 mL fractions were collected (Lobb et al., 2015). Fractions were concentrated using Cytiva Vivaspin (100 kDa MWCO) and protein concentrations were determined by bicinchoninic acid (BCA) assay (ThermoFisher, Waltham, MA). For differential ultracentrifugation (dUC), plasma was centrifuged at 3,000 x g for 5 min (4,000 rpm; Eppendorf 5910R centrifuge) and then at 100,000 x g for 2 h (287,000 rpm; Beckman Coulter SW 60 Ti Swinging Bucket Rotor) to isolate EVs. The EV pellets were washed in PBS and centrifuged again at 100,000 x g for 2 h. Vesicles were analyzed for size, shape, and concentration using the NanoSight NS300 (Malvern Instrument, Westborough, MA) or the Zetaview X30 (Particle Matrix, Mebane, NC) according to the manufacturers’ protocols.

### 2.3 Nanoparticle Tracking Analysis (NTA)

Vesicles were analyzed by NTA for size, shape, and concentration using the NanoSight NS300 (Malvern Instrument, Westborough, MA) or the Zetaview X30 (Particle Matrix, Mebane, NC) according to the protocols by the manufacturers. NanoSight employs laser light scattering and nanoparticle tracking analysis (NTA 2.3 software). Zetaview has the additional capability of measuring EV zeta potential by observing the effect of an electric field on particle movement. Video capture (3 × 30-second captures) of Brownian motion was used to determine the size distribution and concentration of particles in PBS.

### 2.4 Negative-Staining Transmission Electron Microscopy (EM)

All grids were glow-discharged for 45 seconds using the PELCO easiGlow™ system (Ted Pella, Inc.) prior to negative staining. EV samples (10 µl) were loaded onto glow-discharged 100 mesh copper grids (Ted Pella, Inc.) and incubated at room temperature for 2 min. Excess liquid was removed by blotting with filter paper. The samples were fixed with 2% uranyl acetate in distilled water and incubated at room temperature for 2 min. Excess stain was gently blotted off with filter paper. The grids were allowed to air dry for 5 minutes at room temperature before imaging on the Tecnai T-12 transmission electron microscope (ThermoFisher).

### 2.5 Imaging Flow Cytometry

For conventional flow cytometry, EVs were stained with DiD (0.2 mg/ml, 1,1’-Dioctadecyl-3,3,3’,3’-Tetramethylindodicarbocyanine; Invitrogen) and PE-labeled anti-CD63 (H5C6, BD Bioscience) and analyzed on a conventional flow cytometer, Symphony A5 (BD Bioscience). 10,000-20,000 events were collected and analysis was performed with FlowJo software (Tree Star Inc). To detect surface IFNγ, EVs were labeled with anti-IFNγ (B27, BV421-conjugate, BD Biosciences) or isotype control antibodies (MOPC-21, BioLegend). Samples were analyzed on a spectral flow cytometer, Cytek Aurora (Cytek), 10,000-20,000 events were collected and post-collection analysis was performed with FlowJo software (Tree Star Inc). Samples were also analyzed on the Amnis Image Stream imaging flow cytometer (Cytek, Bethesda, MD) with 500,000-800,000 events collected and analyzed by IDEAS® Image Analysis Software (Cytek).

### 2.6 Multiplex EV Surface-protein Markers

EVs isolated from UHPi, UHPp, and IVIg were subjected to immune profiling for 39 known EV surface proteins by flow cytometry using the human MACSPlex EV kit IO (Miltenyi Biotec, Auburn, CA) and detected by flow cytometry (Cytek Aurora, Cytek Biosciences) with fluorescence calibration using Molecular Equivalents of Soluble Fluorophore (MESF) standards (Quantum APC MESF Beads, Bangs Laboratories) as previously described (Nguyen et al., 2024; Welsh et al., 2022). The MACSPlex beads are uniform in size (4.8 µm), barcoded based on fluorescence on the 488 and 561 laser channels, with detection on the 647 nm laser channels. The EV-bead complexes were detected using allophycocyanin (APC)-labeled tetraspanin markers (CD9, CD63, and CD81). Control samples consisting of capture beads alone and capture beads incubated with detection antibodies were acquired at the same time as the samples and used for autofluorescence and nonspecific-binding background fluorescence subtraction. Samples were acquired on the Cytek Aurora flow cytometer and analysis was performed using FlowJo and MPA_PASS_ software as previously described.

EV samples were also analyzed by MILLIPLEX MAP Human Cytokine/Chemokine Magnetic Bead Panel I kit (HCYTOMAG60K, Millipore Sigma, Burlington, MA) that were run on a FlexMAP 3D (Luminex, Austin, TX). Samples were analyzed in both biological and technical triplicates, standard curves were generated for each cytokine, and median fluorescent intensities were transformed into concentrations by five-point, nonlinear regressions (FlowJo software).

### 2.7 Proteinase K Digestion

EVs derived from IVIg by dUC (61.4 µg in 200 µL reaction) were subjected to proteinase K (2 µg/mL) digestion for 1-30 min at 4°C. Reactions were stopped using boiled 0.2 mM phenylmethylsulfonyl fluoride (PMSF) and samples subjected to multiplex cytokine assay by Luminex.

### 2.8 Lipoprotein Depletion

Lipoprotein depletion from SEC-derived IVIg EVs was performed using the LipoMin reagent according to the manufacturer’s protocol (Reliance Biosciences Inc., Taiwan). Briefly, EVs were gently mixed with LipoMin beads and incubated for 10 min at room temp. Lipoprotein-conjugated beads were removed using a Magnetic Separation Rack (Cell Signaling, Danvers, MA).

### 2.9 Cell Culture, EV Treatment, and IFNγ Stimulation

Human epidermoid A431 carcinoma cells were cultured in complete Dulbecco’s Complete Modified Eagle Medium (DMEM) containing 10% FBS (Peak Serum, Fort Collins, CO), L-glutamine, and P/S. Upon reaching 80% confluency, cells were washed with 1xPBS, and switched to serum-free medium (DMEM, L-glutamine, and P/S) and treated with IVIg EVs (0-9 µg/mL) for 24 h. Cells were then stimulated with recombinant IFNγ (3-10 ng/mL; ThermoFisher) for 10-15 min, washed with PBS, and lysed in complete lysis buffer (50 mM Tris-HCl pH 7.5, 150 mM NaCl, 5 mM EDTA, and 1% Triton X-100) supplemented with PMSF, protease and phosphatase inhibitors, and heated to 95°C for 10 min with Laemmli buffer.

### 2.10 Immunoblotting

Proteins were resolved by SDS-PAGE (Bio-Rad Labs, Hercules, CA), transferred to nitrocellulose membranes, blocked for non-specific binding, and immunoblotted. Due to the nature of SEC for EVs, equal volume loading was performed for fractions 1-4 (Void) and 5-9 (EV-rich) while equal protein levels were loaded for fractions 5-9 and 10-14 (protein-rich). Briefly, the combined fractions were incubated with Laemmli buffer with (for human IgG) or without (for CD63) β-mercaptoethanol. Infrared bands were visualized and quantified by the LI-COR Odyssey imaging system (LI-COR Biosciences), ChemiDoc (Bio-Rad Labs, Hercules, CA), or by FluorChem (ProteinSimple, San Jose, CA). Protein signals were quantified using ImageJ and were normalized to corresponding loading control bands (β-actin or GAPDH).

### 2.11 Engineering IFNγ-eEVs

The DNA sequence encoding the fusion protein comprised of the signal peptide (SP), IFNγ coding sequence, cMyc tag, and lactadherin C1C2 domains was synthesized by GenScript (Piscataway, NJ) and inserted into the pCDH-EF1α-MCS-T2A-Neo lentivirus plasmid (System Biosciences, Palo Alto, CA), generating the pCDH-EF1α-hIFNγ-cMyc-C1C2 vector. Plasmid DNA was purified using the Qiagen Maxi Prep Kit (ThermoFisher). HEK293T/17 cells were cotransfected with the following plasmids pCDH-EF1α-hIFNγ-cMyc-C1C2, pRSV-Rev, PMDLg/pRRE, and pMD2.g (Addgene, Watertown, MA) using Lipofectamine 3000 (ThermoFisher) according to the manufacturer’s protocol. Lentivirus was purified from the conditioned medium using PEG-8000 (Sigma-Aldrich). HEK293T/17 cells were transduced with the concentrated lentivirus in the presence of 0.8 ug/mL polybrene and selected with G418 antibiotics. Construct sequence will be made available upon request.

### 2.12 Cold Ethanol Precipitation

SEC- or dUC-isolated eEVs were precipitated using ice-cold ethanol (100%; 9:1 volume), incubated at -80°C for 2 h, centrifuged at 13,000 x g for 30 min at 4°C, and washed with ethanol (90%). The EV pellets were air-dried and resuspended in PBS. Alternatively, conditioned medium was first concentrated and then subjected to cold ethanol precipitation prior to SEC purification. The resuspended pellet was loaded over SEC columns and EV fractions were combined and concentrated.

### 2.13 Statistics

Results are reported as mean ± standard error of the mean (SEM) utilizing both technical and biological triplicates. 2-tailed Student’s *t*-test was performed when appropriate. For the Luminex data, each sample was also performed in technical triplicates. Analyses and graph creation were conducted using GraphPad Prism v10. Significance was defined as follows: ns P > 0.05; *P≤0.05; P-value: * P ≤ 0.05; ** P ≤ 0.01; *** P ≤ 0.001; **** P ≤ 0.0001.

## 3 Results

### 3.1 EVs Derived from IVIg and SCIg

To establish the presence of EVs, we performed SEC on IVIg, SCIg, individual UHPi, and pooled UHPp. Due to limited SCIg availability, most data derive from IVIg, with SCIg included where feasible. Plasma (1 mL) samples were loaded over SEC columns, yielding 15 fractions (1 mL each). Protein levels, measured by BCA assay, rose rapidly in fractions 11-15 (Figure 1A). Western blotting detected IgG beginning in fractions 8-9, with a sharp increase from fractions 10-15 (Figure 1B). Based on the manufacturer’s protocol, we assigned fractions 1-4 as the column void volume, fractions 5-9 as EV-rich, and fractions 10-14 as protein-rich (Figure 1A). Fractions 5-9 were pooled and concentrated. Nanoparticle tracking analysis (NTA) revealed the presence of particles in all samples (Figure 1C). NTA measured larger particle sizes in IVIg and SCIg compared to UHPi and UHPp (Figure 1D; Table 1) but with similar vesicle concentrations across plasma samples (Figure 1E; Table 1). TEM confirmed the presence of intact nanometer-sized, membrane-bound particles (Figure 1F; Figure S1). To our knowledge, this is the first demonstration of EVs in IVIg and SCIg.

**FIGURE 1.**
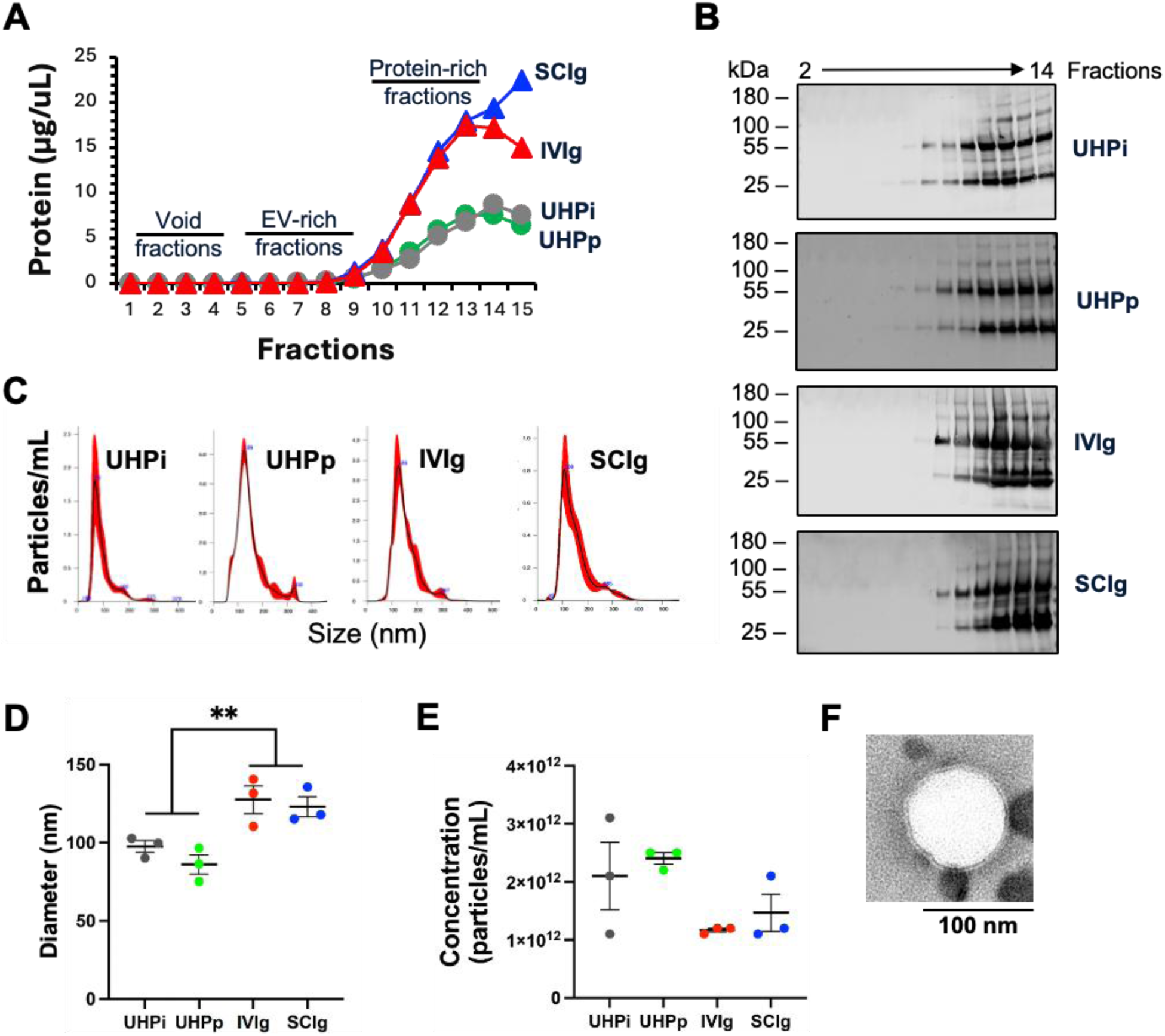
Isolation and characterization of EVs in IVIg and SCIg. (A) EVs were isolated from UHPi (gray), UHPp (green), IVIg (red), and SCIg (blue) by SEC. Fifteen fractions (1 mL) were collected, and protein concentrations were determined by BCA assay. Fractions: 1-4 (void); 5-9 (EV-rich), and 10-15 (protein-rich). (B) Immunoblotting of each fraction with anti-human IgG antibodies. (C-E) NTA profile of pooled, concentrated EV-rich fractions (C), diameter (D), and concentration (particles/mL input volume) (E). (F) Representative TEM image of IVIg EVs showing lipid bilayer. Size bar = 100 nm. P-value: * P ≤ 0.05. BCA, bicinchoninic acid assay; EV, extracellular vesicles; IgG, immunoglobulin; IVIg, intravenous immunoglobulin; NTA, nanoparticle tracking analysis; SCIg, subcutaneous immunoglobulin; SEC, size-exclusion chromatography; TEM, transmission electron microscopy; UHPi, individual unprocessed human plasma; UHPp, pooled unprocessed human plasma; n=3.

**TABLE 1.**

|  | **UHPi** |  | **UHPp** |  | **IVIg** |  | **SCIg** |  |
| --- | --- | --- | --- | --- | --- | --- | --- | --- |
|  | **Concentration**<br>**(Particles/mL)** | **Size**<br>**(nm)** | **Concentration**<br>**(Particles/mL)** | **Size**<br>**(nm)** | **Concentration**<br>**(Particles/mL)** | **Size**<br>**(nm)** | **Concentration**<br>**(Particles/mL)** | **Size**<br>**(nm)** |
|  | 2.50E+12 | 102.7 | 3.10E+12 | 86.3 | 1.10E+12 | 110.3 | 1.10E+12 | 135.6 |
|  | 2.50E+12 | 99.7 | 2.10E+12 | 96.4 | 1.20E+12 | 140.4 | 1.20E+12 | 117.8 |
|  | 2.20E+12 | 90.1 | 1.10E+12 | 75 | 1.20E+12 | 131.6 | 2.10E+12 | 115.1 |
| **Mean** | **2.40E+12** | **97.5** | **2.10E+12** | **85.9** | **1.17E+12** | **127.4** | **1.47E+12** | **122.8** |
| **SEM** | **1.00E+11** | **3.8** | **5.77E+11** | **6.2** | **3.33E+10** | **8.9** | **3.18E+11** | **6.4** |
| **P-value** | **UHPi vs IVIg/SCIg** |  |  |  | **0.0003** | **0.0368** | **0.0488** | **0.0275** |
| **P-value** |  |  | **UHPp vs IVIg/SCIg** |  | **0.1818** | **0.0187** | **0.3910** | **0.0144** |

IVIg, intravenous immunoglobulin; SCIg, subcutaneous immunoglobulin; UHPi, individual unprocessed human plasma; UHPp, pooled unprocessed human plasma

### 3.2 CD63-positive EVs in IVIg

To confirm the presence of EVs we used the MACSPlex EV immuno-oncology Kit which detects 37 different surface epitopes with two isotype controls. We profiled 37 surface proteins, including the tetraspanins CD9, CD63 and CD81 (Figure 2A). CD9, CD63 and CD81 were detected in UHPi EVs while IVIg EVs displayed higher CD63 but markedly lower CD9 and CD81 (Figure 2A, B). Consistent with prior reports, UHPi EVs were enriched in platelet markers (CD42b, CD42a, CD62P; Figure 2B) and the Human Leukocyte Antigen Family (HLA) proteins (HLA-ABC, HLA-DR/DP/DQ; Figure 2B), whereas these were markedly reduced in IVIg (Cappellano et al., 2021; Eustes and Dayal, 2022). Conversely, IVIg EVs were strikingly enriched in B- and T-cell markers (CD19, CD20, and CD69), along with stemness markers, (CD29, CD133/1, CD326, and ROR1; Figure 2B). Thus, UHPi EVs derive largely from platelets, while IVIg EVs reflect greater B-, T-, and stem cell origins.

**FIGURE 2.**
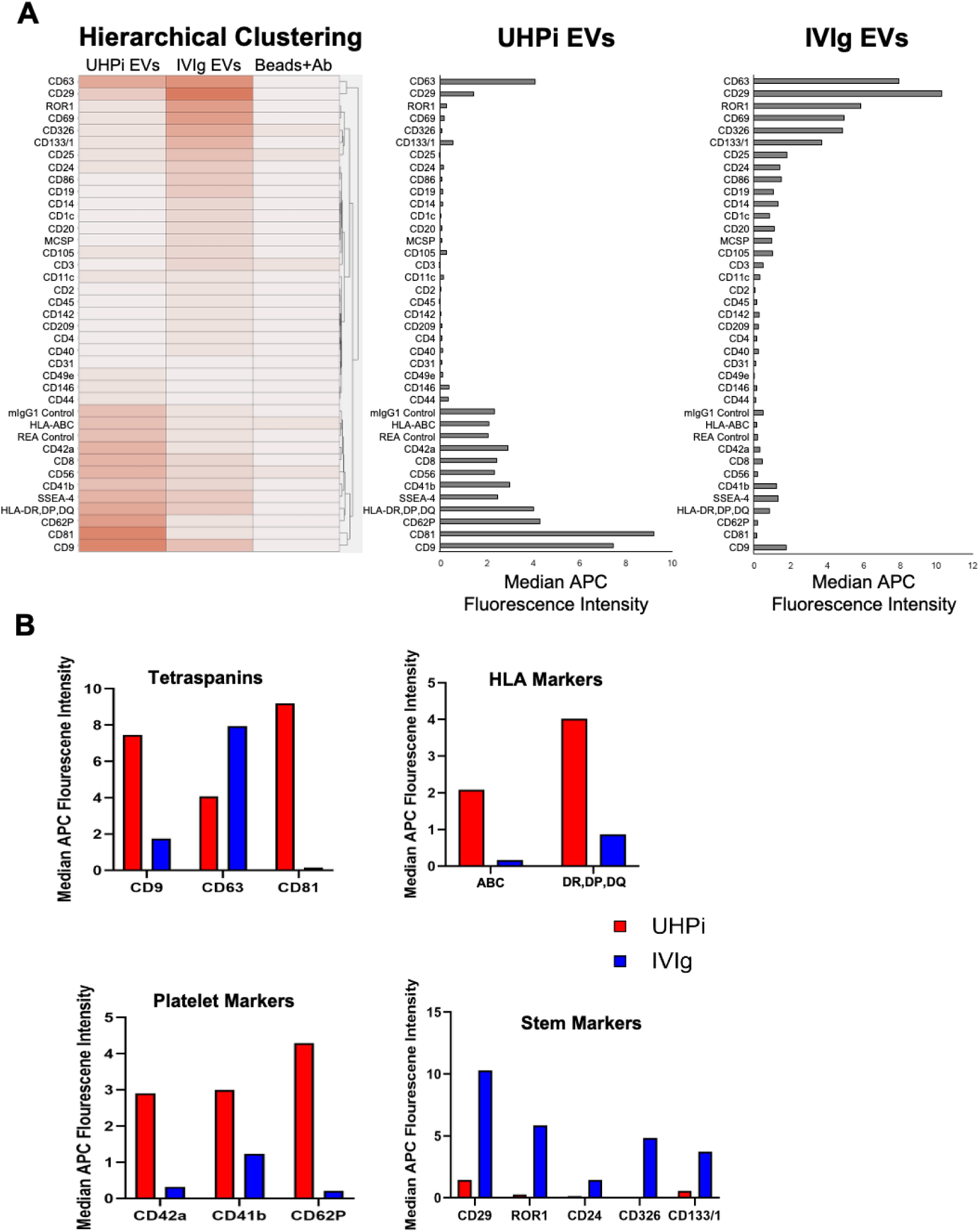
Multiplex bead-based EV flow cytometry assay for surface markers. EVs enriched from UHPi and IVIg by SEC and dUC. Surface marker profiles were measured using Miltenyi MACSPlex EV kit IO with MESF-calibrated flow cytometry. (A) Heatmap and hierarchical clustering of MACSPlex markers in UHPi EVs, IVIg EVs, and bead/antibody controls (left). Median APC intensity (MESF, background-subtracted) of mixed tetraspanin antibodies (CD9/CD81/CD63) on EVs captured by 39 marker beads (middle/right). (B) Enlarged values for the tetraspanins (CD9, CD63, and CD81), HLA markers (HLA-ABC and HLA-DR, DP,DQ), platelet markers (CD42a, CD41b, and CD62p), and stemness markers (CD29, ROR1, CD24, CD326, CD133/1). dUC, differential ultracentrifugation; EV, extracellular vesicles; Human Leukocyte Antigens; HLA; IVIg, intravenous immunoglobulin; MESF, Molecular Equivalents of Soluble Fluorophore; SEC, size-exclusion chromatography; UHPi, individual unprocessed human plasma.

Next, using the Cytex Amnis imaging flow cytometry high-resolution images of individual particles, we analyzed dUC- and SEC-isolated UHPp and IVIg EVs. Representative composite images showed individual EVs in bright field (BF), PE (CD63), APC (DiD), and side scatter (SSC) channels (Figure 3A). Gating density plots revealed similar percentages of potential EVs by size (Figure 3B, left panel, scatter imaging, gate R1;), and fluorescence imaging (Figure 2B, right two panels, gate R2) in DiD^+^ control UHPp and CD63^+^ EVs. Gating revealed similar percentages of putative EVs (gate R1) in UHPp and IVIg by both methods (Figure 3B-C, left). However, DiD^+^CD63^+^ EVs were more abundant in IVIg than UHPp, independent of isolation method (Figure 3C, right). These findings confirm CD63^+^ EVs in IVIg, consistent across donor-pooled lots and isolation approaches.

**FIGURE 3.**
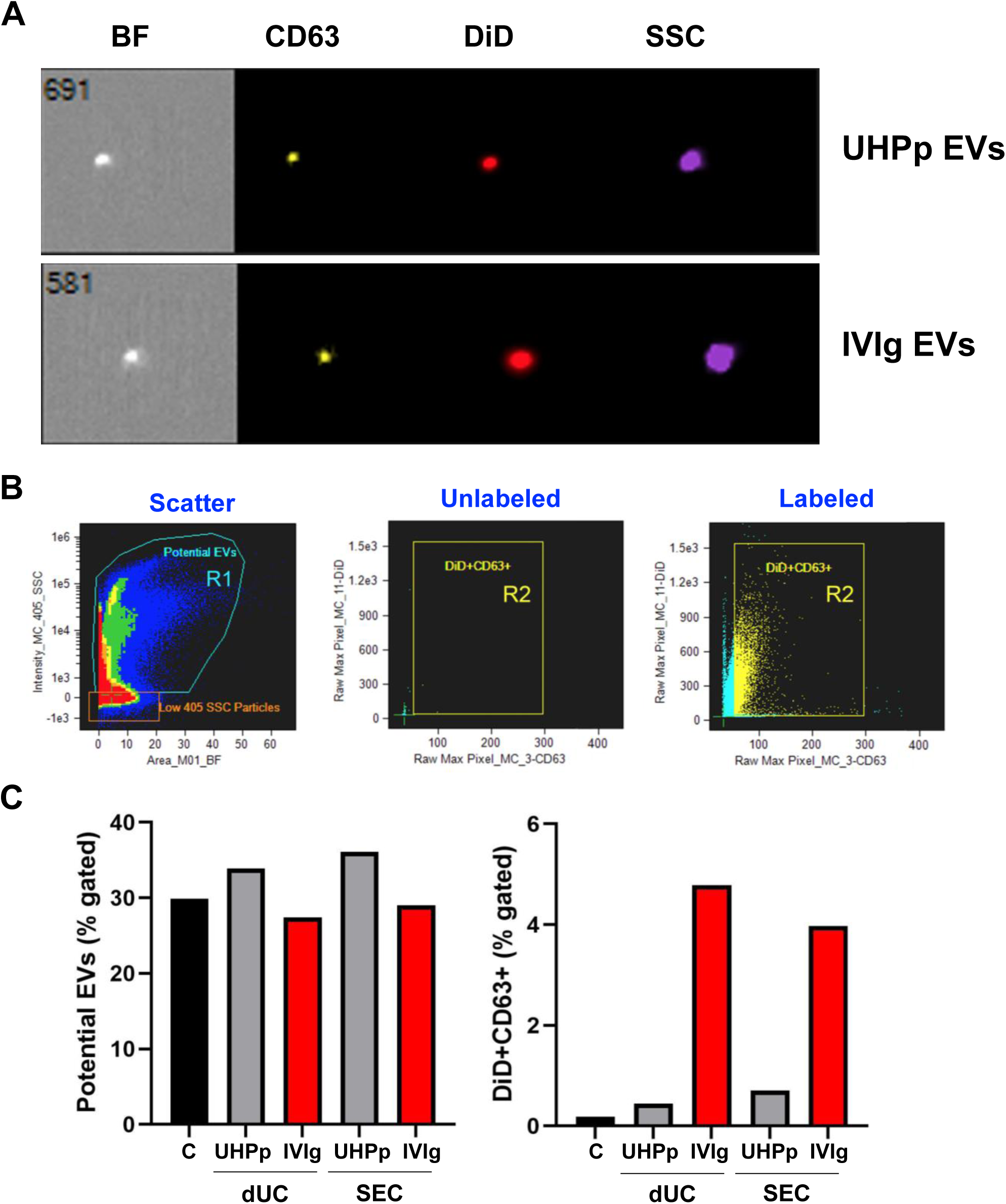
Flow cytometry phenotyping of CD63-positive EVs. UHPp, and IVIg EVs isolated using dUC or SEC were labeled with DiD and CD63-PE and analyzed by imaging flow cytometry. (A) Representative images of UHPp and IVIg EVs show morphology, BF, CD63, DiD, and scatter channels. (B) Scatter was used to gate out debris (left, gate R1). Fluorescent dot plots of unlabeled samples show background signal (middle) and labeled samples identified DiD^+^ and CD63^+^ events (right, gate R2). (C) Summary plots of the frequency of EVs identified by scatter (gate R1, left) or fluorescence (gate R2, right) of UHPp, and IVIg EVs isolated using dUC or SEC. BF, bright field; DiD, 1,1′-dioctadecyl-3,3,3′,3′- tetramethylindodicarbocyanine, 4-chlorobenzenesulfonate salt; dUC, differential ultracentrifugation; EV, extracellular vesicles; IVIg, intravenous immunoglobulin; PE, phycoerythrin; SEC, size-exclusion chromatography; UHPp, pooled unprocessed human plasma. Note: two different lots of UHPp and IVIg were used for these experiments.

Finally, CD63 levels were assessed by immunoblotting across SEC fractions from UHPi, IVIg, and SCIg (Figure S2) (Welsh et al., 2024a). Due to drastic differences in protein concentration, equal volumes of column void and EV-rich fractions were loaded, while equal protein concentrations were used for the EV-rich and protein-rich fractions. CD63 appeared only in the EV-rich fractions but were undetectable in the column void or protein-rich fractions (Figure S2). IgG heavy (55kDa) and light (25 kDa) chains were present in both the EV-rich and protein-rich lanes. Furthermore, conventional flow cytometry of the membrane lipid dye DiD-stained EVs with PE-anti-CD63 showed most DiD^+^ IVIg EVs were CD63^+^ (Figure S3).

In summary, using three different flow cytometry methods and immunoblotting, we have convincingly shown the presence of CD63 in IVIg EVs.

### 3.3 IVIg EVs are Associated with Cytokines, Chemokines, and Growth Factors

Cytokines modulate immunity and inflammation (Turner et al., 2014), circulating freely, residing inside EVs, or binding EV surfaces through the biomolecular corona (Fitzgerald et al., 2018; Jung et al., 2020). Given IVIg lineage differences compared to UHP, we explored associated cytokines to functional provide functional insights. Magnetic bead-based assays of SEC fractions from IVIg, UHPi and SCIg revealed distinct cytokine profiles by source and EV- versus protein-rich fractions (Figure 4A). RANTES, a chemokine that plays a crucial role in immune regulation, particularly in the recruitment of T cells, monocytes, and other immune cells to sites of acute and often chronic inflammation (Barczak et al., 2023), was expressed up to 1000-fold higher levels in UHPi (across both fractions) compared to IVIg/SCIg. Another pro-inflammatory cytokine, IL-18, was elevated in the UHPi protein-rich fractions, with levels rising after fraction 10. The potent anti-inflammatory cytokines IL-4 and IL-10 were higher in IVIg/SCIg (IL-4 detected in early EV-rich fractions, IL-10 mainly in the protein-rich fractions). Dual function IL-3 and IFNγ were elevated in IVIg/SCIg, with IFNγ uniquely showing a bell-shaped distribution spanning the EV/protein interface. In summary, differential EV-rich and protein-rich fraction cytokine distribution patterns across UHPi, IVIg, and SCIg were revealed (Figure 4A; Figure S4), likely indicating unique therapeutic functions and EV lineage origins of IVIg (and, by extension SCIg) compared with UHPi EVs.

**FIGURE 4.**
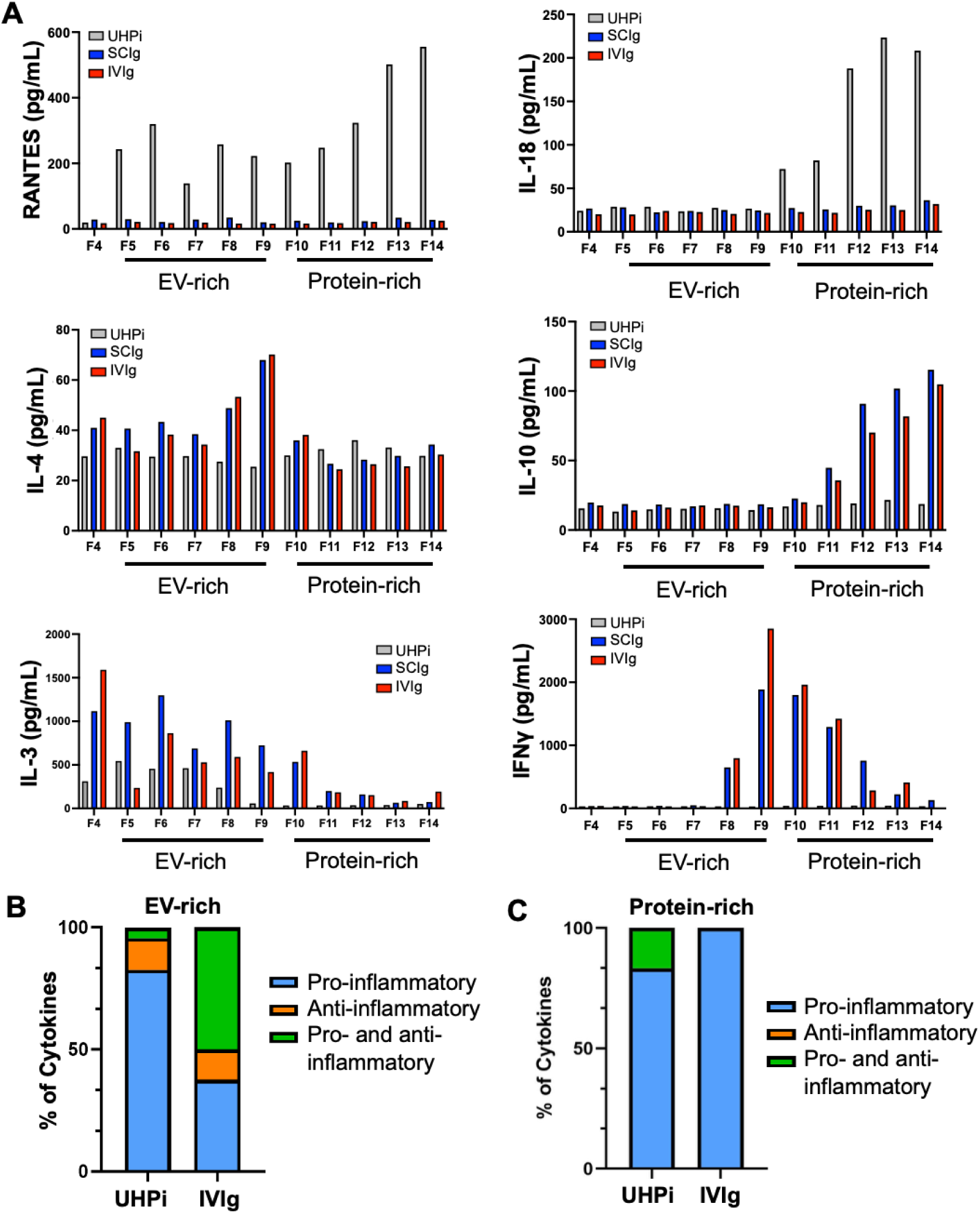
Comparison of surface‒associated cytokines in UHPi, IVIg, and SCIg SEC fractions. (A) Representative fractions 4-14 from UHPi (gray), IVIg (red), and SCIg (blue) purified over SEC and subjected to cytokine analysis by Luminex. Values were plotted and represented as bar graphs for each fraction. The lines under each graph demarcate the EV- and protein-rich fractions. Individual representative plots of pro-inflammatory cytokines RANTES and IL-18, anti-inflammatory cytokines IL-4 and IL-10, and dual pro-/anti-inflammatory cytokines IL-3, and IFNγ. (B-C) Bar graphs showing differences in pro- (blue), anti- (orange), or both pro- and anti-inflammatory (green) cytokines in the EV-rich (B) and protein rich (C) fractions of UHPi and IVIg. EV, extracellular vesicles; IFNγ, interferon gamma; IL, interleukin; IVIg, intravenous immunoglobulin; RANTES, regulated upon activation normal T cell expressed and secreted; SEC, size-exclusion chromatography; SCIg, subcutaneous immunoglobulin; UHPi, individual unprocessed human plasma. UHPi, n=6; IVIg, n=3 different lots.

To validate and expand these observations, EV-rich and protein-rich fractions were pooled from 6 UHPi and 3 additional IVIg lots (each pooled from >1000 donors) and analyzed by Luminex cytokine profiling. Results showed IVIg EV-rich fractions contained significantly lower levels of 22 pro-inflammatory cytokines than UHPi-derived EVs but higher levels of 8 other cytokines (Figure S5; Table 2). In EV-rich fractions, UHPi carried more pro-inflammatory cytokines (Figure 4B, blue) while IVIg was enriched in dual pro- and anti-inflammatory species (Figure 4B, green). In the protein-rich fractions, both UHPi and IVIg carried more pro-inflammatory cytokines (Figure 4C, blue). Cytokine profiles were similar between 3 independent lots of UHPp and 3 UHPi EVs (Figure S6), with only IL-22 and PDGF-AA/BB displaying significantly reduced levels and 4 cytokines below the detection limit. SCIg was compared to UHPi independently, owing to limited SCIg availability and assay performance timing. SCIg EVs resembled IVIg EVs showing lower RANTES and higher IFNγ (Figure S7). Finally, IVIg protein-rich fractions showed lower levels of 12 pro-inflammatory cytokines (including RANTES) and higher levels of 7 others (including IL-10) than UHPi (Figure S8). Thus, IVIg processing enriches EVs and soluble cytokines associated with reduced inflammation.

**TABLE 2.** Fold change in the levels of cytokines, chemokines, and growth factors detected by the Luminex multiplex assay in the EV-rich fractions of IVIg compared to UHPi.

| EV-rich |  |  |  |  |
| --- | --- | --- | --- | --- |
|  | UHPi | IVIg |  |  |
| Cytokines | AVE±SEM (pg/mL) | AVE±SEM (pg/mL) | Fold change | p-value |
| RANTES | 12023±4368 | 19.22±0.67 | 0.001599 | 0.0238 |
| sCD40L | 80.11±22.51 | 22.67±1.17 | 0.282986 | 0.0238 |
| IL-15 | 75.00±20.87 | 21.56±1.06 | 0.287467 | 0.0238 |
| PDGF-AB/BB | 113.9±5.45 | 38.67±6.74 | 0.339508 | 0.0357 |
| FGF-2 | 45.61±7.38 | 18.33±0.88 | 0.401886 | 0.0238 |
| PDGF-AA | 70.63±6.88 | 32.72±1.01 | 0.463259 | 0.0357 |
| TGFα | 47.92±7.01 | 22.72±0.06 | 0.474124 | 0.0238 |
| IL-18 | 49.75±3.85 | 23.72±0.75 | 0.476784 | 0.0238 |
| Eotaxin | 30.17±3.34 | 14.83±0.10 | 0.491548 | 0.0238 |
| IL-17A | 57.73±9.28 | 27.61±1.28 | 0.47826 | 0.0357 |
| IL-17F | 45.92±7.09 | 22.78±0.95 | 0.49608 | 0.0238 |
| TNFα | 40.06±5.25 | 21.39±0.64 | 0.533949 | 0.0238 |
| TNFβ | 41.33±3.90 | 22.17±1.17 | 0.536414 | 0.0357 |
| MIP-1β | 27±3.60 | 14.44±0.59 | 0.534815 | 0.0238 |
| IL-17E | 50.03±3.46 | 27.11±0.29 | 0.541875 | 0.0238 |
| Fractalkine | 22.22±3.48 | 12.56±0.11 | 0.565257 | 0.0238 |
| GROα | 31.03±2.66 | 17.67±1.07 | 0.569449 | 0.0238 |
| IL-1β | 50.92±5.45 | 30.72±0.89 | 0.603299 | 0.0238 |
| IL-2 | 39.56±2.18 | 27.44±1.04 | 0.69363 | 0.0238 |
| VEGF-A | 55.17±7.01 | 41.28±2.71 | 0.748233 | 0.0476 |
| IL-8 | 21.67±1.56 | 16.5±0.76 | 0.761421 | 0.0238 |
| IL-6 | 23.75±1.46 | 18.71±0.56 | 0.787789 | 0.0476 |
| IL-3 | 35.58±8.69 | 715.3±93.65 | 20.10399 | 0.0238 |
| IL-7 | 63.81±11.02 | 649.2±110.4 | 10.17395 | 0.0238 |
| MIP-1α | 70.36±17.27 | 376.7±26.8 | 5.353894 | 0.0238 |
| MDC | 40.75±3.60 | 216.5±16.31 | 5.312883 | 0.0238 |
| IFNγ | 50.22±5.63 | 233.3±83.75 | 4.64556 | 0.0238 |
| GM-CSF | 413.2±122.5 | 1250±224.2 | 3.025169 | 0.0238 |
| MCP-3 | 36.44±4.96 | 63.67±1.53 | 1.747256 | 0.0238 |
| MIG | 29.14±3.32 | 44.33±3.42 | 1.521277 | 0.0476 |

IVIg, intravenous immunoglobulin (n=3); UHPi, individual unprocessed human plasma (n=6)

### 3.4 IFNγ is Associated on the Surface of IVIg EVs

The anti-inflammatory effects of IVIg have been suggested to be mediated through the suppression of IFNγ-mediated IFNGR/JAK/STAT1 activation (Park-Min et al., 2007). Here, we detected high levels of IFNγ in the IVIg EV-rich fractions. To determine whether IFNγ is surface-bound or encapsulated, dUC-isolated IVIg EVs were subjected to proteinase K (PK) digestion. PK-treated EVs were then subjected to cytokine analysis by Luminex which revealed reduced IFNγ levels by ∼75% without affecting MIP-1β (Figure 5A), suggesting that approximately 25% of IFNγ is encapsulated within the IVIg EVs. The Luminex and PK results were further validated by single EV flow cytometry which measured higher IFNγ in IVIg and SCIg EV-rich fractions than UHPi, and in EV-rich versus protein-rich fractions (Figure 5B-C). Since IgG binds to EV membranes and IFNγ-specific antibodies were present, immunoblotting demonstrated that these antibodies recognize both monomeric (14 kDa) and dimeric (28 kDa) IFNγ. Like previous findings (Lunn et al., 1992; Zlateva et al., 1999), we demonstrate that IFNγ can persist as dimers even under reducing conditions (Figure S9). Thus, IFNγ likely associates with IVIg EVs through binding to IFNγ-specific antibodies.

**FIGURE 5.**
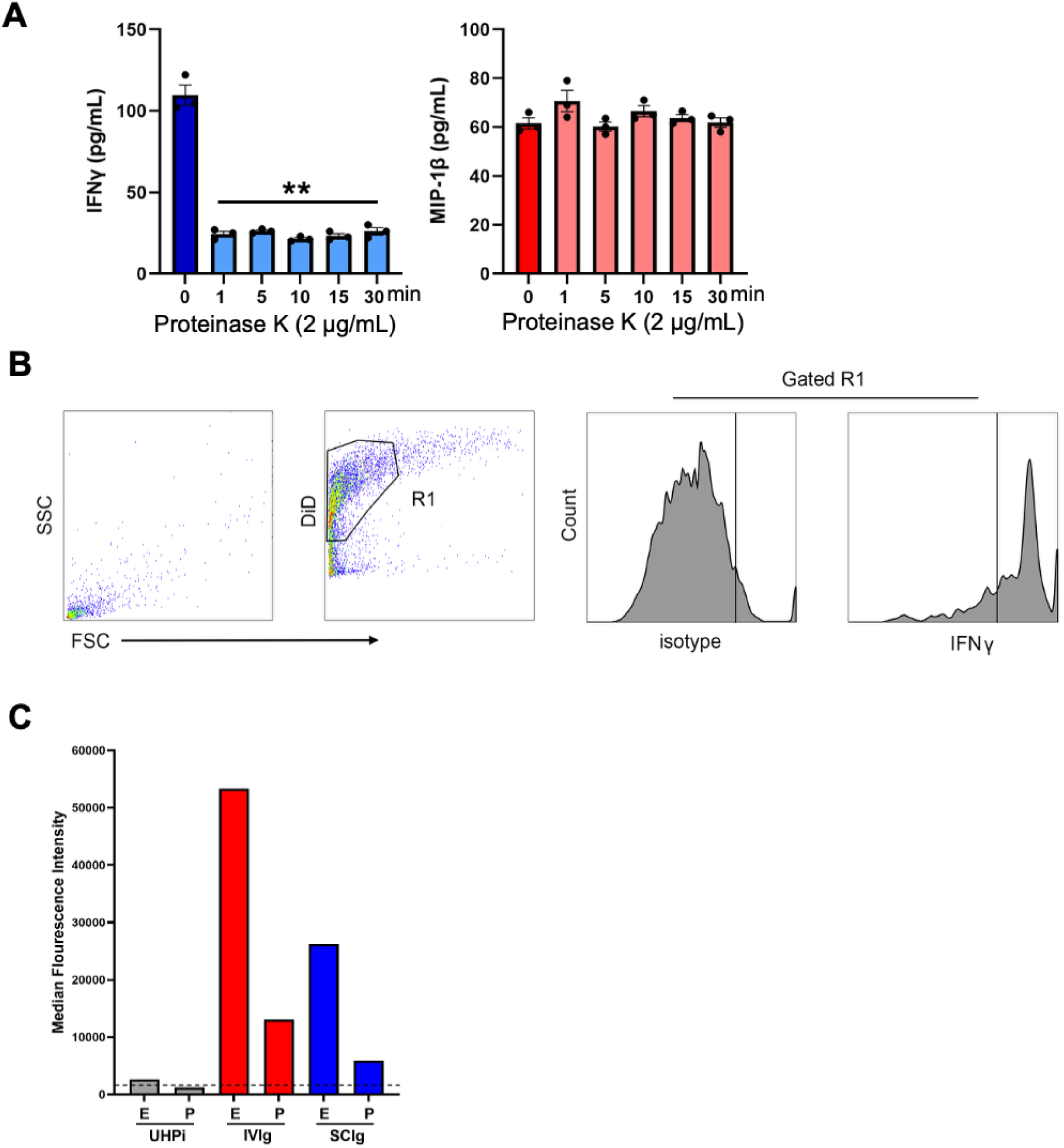
IVIg EVs contain high levels of surface-associated IFNγ. (A) IVIg EV-rich fractions from SEC were subjected to proteinase K digestion for 1-30 min and levels of IFNγ and MIP-1β were analyzed by Luminex. (B) Flow cytometry analysis of different enrichment fractions. Particles were isolated from UHPi, SCIg, and IVIg by SEC. An EV gate was established based on size and lipophilic dye labeling with DiD. (C) Histograms depict representative fluorescence levels following labeling with an IFNγ antibody or isotype control. Summary plot contains MFI values of IFNγ (bars) or isotype (dashed line) staining of various fractions (E, EV-rich; P, protein-rich) isolated by dUC or SEC. Bars outline fractions containing EVs. DiD, 1,1′-dioctadecyl-3,3,3′,3′-tetramethylindodicarbocyanine; 4-chlorobenzenesulfonate salt; dUC, differential ultracentrifugation; EVs, extracellular vesicles; IFNγ, interferon gamma; IVIg, intravenous immunoglobulin; MFI, median fluorescence intensity; MIP-1β, macrophage inflammatory protein-1 beta; SEC, size-exclusion chromatography; UHPi, individual unprocessed human plasma.

### 3.5 IVIg EVs Inhibit IFNγ-induced JAK/STAT1 Activation

To determine whether EVs play a role in the IVIg inhibitory effect of the IFNGR/JAK/STAT1 signaling pathway, we used A431 cells, which possess an intact and activatable IFNGR/JAK/STAT1 signaling pathway. Recombinant IFNγ (10 ng/mL) induced STAT1 (p-STAT1 phosphorylation (p-STAT1) in A431 cells after 5 min, peaking at 10-15 min (Figure 6A). To isolate larger EV quantities required for the IVIg inhibition experiments, EVs were isolated from 10 mL of 2 independent lots of IVIg plasma by SEC, collecting 5 mL fractions each (Figure S10A). A sharp rise in protein levels was observed in fractions 6 and 7. Immunoblotting confirmed RANTES in UHPi but not IVIg (Figure S10B). Surprisingly, these EVs did not activate p-STAT1, despite substantial bound IFNγ (Figure 6B). Pre-incubation with IVIg fraction 4 (0-9 µg/mL; 24 h) dose-dependently inhibited subsequent recombinant IFNγ-induced (10 ng/mL; 15 min) p-STAT1, displaying an inverse correlation between p-STAT1 levels and IVIg EV dose in response to IFNγ stimulation on immunoblots (Figure 6C). Signals were quantitated from three independent experiments (biological replicates) and plotted (Figure 6C, right panel). The experiment was repeated using a lower concentration of IFNγ (3 ng/mL) and a single concentration of IVIg EVs (9 µg/mL) which resulted in dramatic p-STAT1inhibition (Figure 6D). Inhibition varied across three different IVIg lots (Figure S10C), but significantly was absent with UHPi/UHPp EVs, even at high doses (Figure S10D).

**FIGURE 6.**
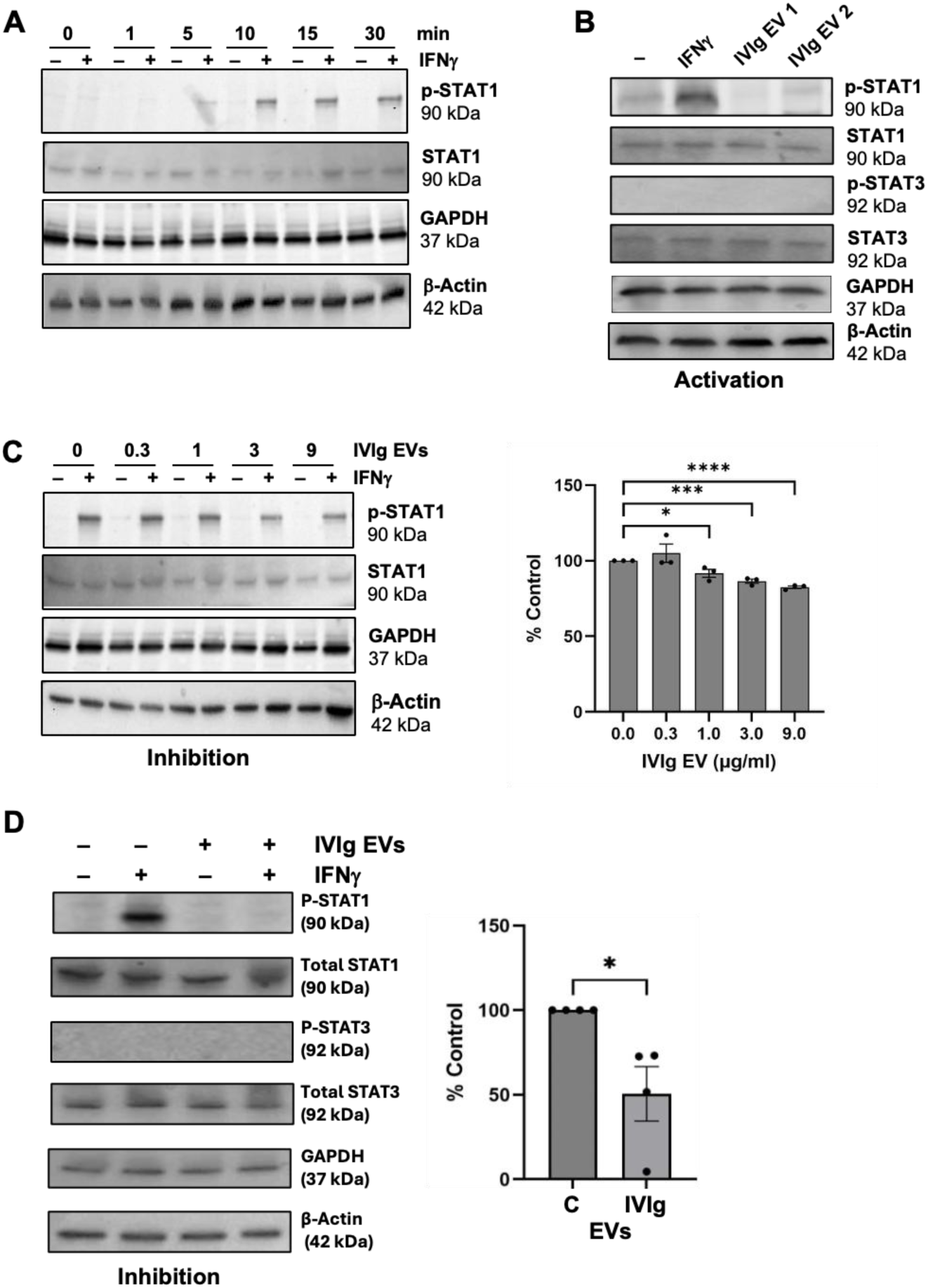
IVIg EVs inhibit IFNγ-mediated JAK/STAT1 activation. (A) Cells cultured in serum-free medium for 24 h were treated with recombinant IFNγ (10 ng/mL) for 1-30 min. Cells were lysed and proteins were immunoblotted for p-STAT1, STAT1, GAPDH and β-Actin. (B) Cells were treated with IFNγ (3 ng/mL), and two preps of IVIg EVs (9 µg/mL) for 15 min and then lysed for immunoblotting for p-STAT1, STAT1, p-STAT3, STAT3, GAPDH and β-Actin. (C) Cells were pre-incubated with IVIg EVs (0-9 µg/mL) for 24 h, treated with IFNγ (10 ng/mL) for 15 min, and lysed for immunoblotting for p-STAT1, STAT1 (loading control), GAPDH and β-Actin. Band intensity was quantitated by ImageJ and inhibition at 9 µg/mL are represented as percent control (n=3; biological replicates) and represented in the graph to the right. (D) Cells were preincubated with IVIg EVs (0-9 µg/mL) for 24 h, treated with IFNγ (3 ng/mL) for 15 min, and lysed for immunoblotting for p-STAT1, STAT1, p-STAT3, STAT3, GAPDH and β-Actin. Band intensity was quantitated by ImageJ and inhibition is represented as percent control (n=4; biological replicates) and represented in the graph to the right. P-value: * P ≤ 0.05; ** P ≤ 0.01; *** P ≤ 0.001; **** P ≤ 0.0001. EV, extracellular vesicles; GAPDH, glyceraldehyde-3-phosphate dehydrogenase; IFNγ, interferon gamma; IVIg, intravenous immunoglobulin; p-STAT1, phosphorylated signal transducer and activator of transcription 1; p-STAT3, phosphorylated signal transducer and activator of transcription 3; STAT1, signal transducer and activator of transcription 1; STAT3, signal transducer and activator of transcription 3.

Finally, to assess whether the inhibitory effects of IVIg on JAK/STAT1 signaling are mediated by non-EV components, Western blotting for lipoproteins was performed. Lipoproteins ApoA1 and ApoB were detected in UHPp but not IVIg EV-rich fractions (Figure S11A). Furthermore, when IVIg EVs were treated with LipoMin, the lipoprotein-depleted EVs neither activated (Supplemental Figure 11B) nor altered the inhibition (Figure S11C) of IFNγ signaling, confirming the effect is EV-mediated and not lipoprotein-mediated.

### 3.6 Dual Inflammatory Roles of IFNγ

To test for dual IFNγ-EV effects, we engineered surface-bound IFNγ on EVs using lentivirus stable cell lines overexpressing a fusion protein comprised of human IFNγ, cMyc, and the lactadherin C-terminal C1 and C2 domains (IFNγ-eEVs; Figure 7A). Lactadherin C1C2 binds phospholipids, such as phosphatidylserine, that are enriched on EV membranes. SEC-isolated EV-rich fractions from transformed cell-conditioned medium (Figure 7B) showed high IFNγ (Figure 7C, early fractions 2-5), which displayed an ∼60 kDa band consistent with the IFNγ-lactadherin fusion protein Figure 7D) employing 2 different antibodies (Luminex and immunoblotting). IFNγ-eEVs activated JAK/STAT1 signaling similar to recombinant IFNγ (Figure 7E) but, after 24 h pre-treatment, suppressed response to recombinant IFNγ (Figure 7F).

**FIGURE 7.**
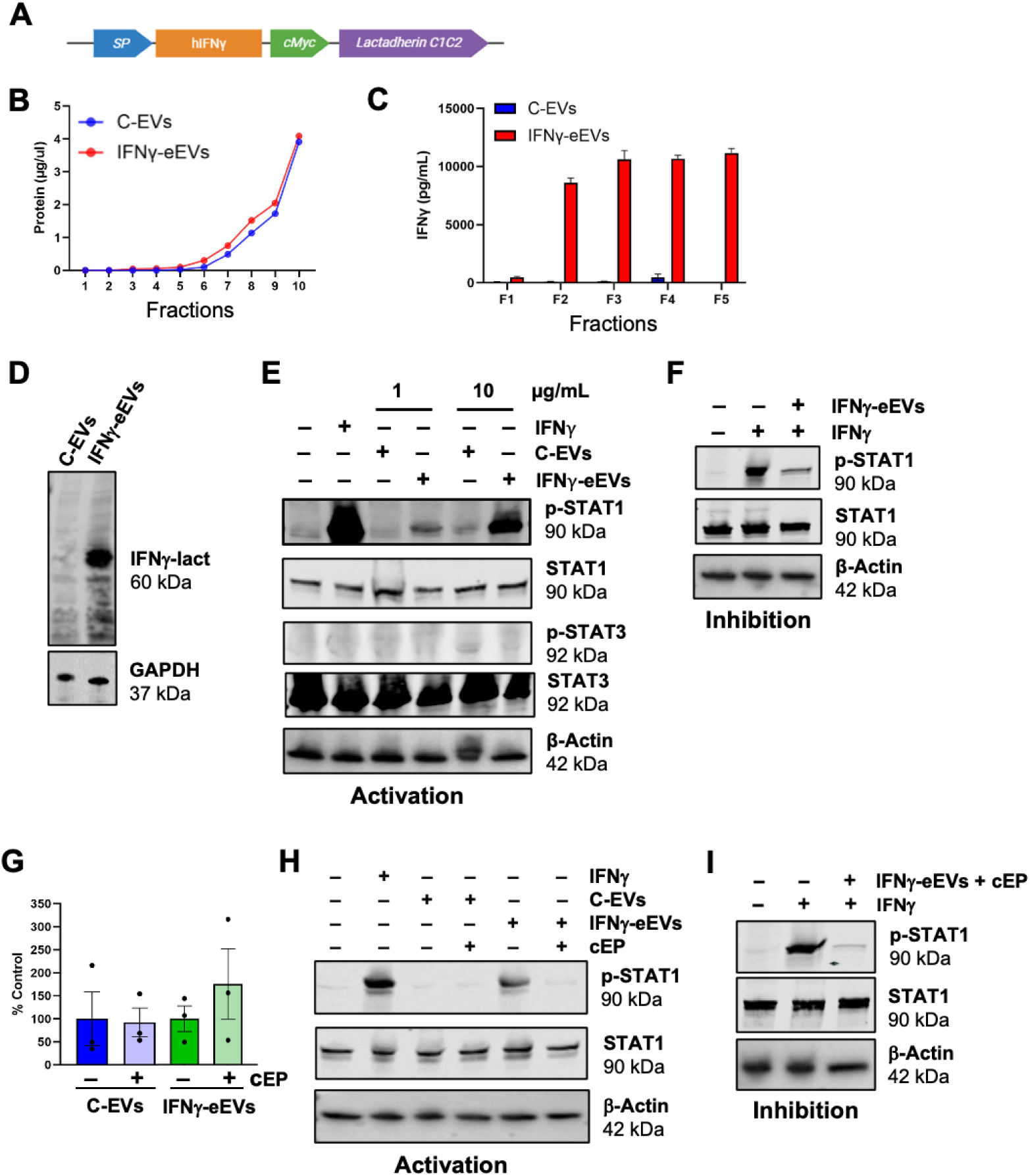
Inactivation of IFNγ by cold ethanol precipitation. (A) Schematic diagram of lentiviral plasmid sequence encoding the IFNγ-cMyc-Lactadherin C1C2 fusion protein (IFNγ-lact). (B) Stable-expressing HEKT/17 cell lines were established and EVs were isolated from conditioned medium by SEC using Izon qEV10 columns. Ten fractions (5 mL) were collected, and protein concentrations were determined by BCA. (C) ELISA by Luminex for IFNγ in the early EV-rich fractions 2-5. (D) Fractions 2-5 of C-EVs and IFNγ-eEVs were pooled, concentrated, and immunoblotted for IFNγ (60 kDa fusion protein). (E) Cells were stimulated with recombinant IFNγ (3 ng/mL), C-EVs (1 and 10 µg/mL) and IFNγ-eEVs (1 and 10 µg/mL) for 15 min. Proteins were lysed and immunoblotted for p-STAT1, STAT1, p-STAT3, STAT3, and β-Actin. (F) Western blot for p-STAT1, STAT1, and β-Actin of cells pre-treated with IFNγ-eEVs (10 µg/mL) for 24 h and were then stimulated with recombinant IFNγ (3 ng/mL) for 15 min. (G) NTA analysis of C-EVs and IFNγ-eEVs before and after cEP expressed as percent control. (H) Cells were treated with recombinant IFNγ (3 ng/mL) or C-EVs and IFNγ-EVs (±EP; 10 µg/mL) for 15 min. Cells were lysed and immunoblotted for p-STAT1, STAT1, and β-Actin. (I) Cells were pre-treated with IFNγ-eEVs (10 µg/mL each) for 24 h and then stimulated with recombinant IFNγ (3 ng/mL) for 15 min. Cells were lysed and proteins immunoblotted for p-STAT1, STAT1, and β-Actin. BCA, bicinchoninic acid assay; C-EVs, control EVs; cEP, cold ethanol precipitation; EV, extracellular vesicles; IFNγ, interferon gamma; IFNγ-eEVs, EVs with surface associated IFNγ-cMyc-Lactadherin; NTA, nanoparticle tracking analysis; p-STAT1, phosphorylated signal transducer and activator of transcription 1; p-STAT3, phosphorylated signal transducer and activator of transcription 3; STAT1, signal transducer and activator of transcription 1; STAT3, signal transducer and activator of transcription 3; STAT3, signal transducer and activator of transcription 3.

Cold ethanol precipitation (cEP), a critical IVIg processing step (Afonso and João, 2016; Belmonte et al., 2025) preserved vesicles numbers (Figure 7G) but blocked IFNγ-eEVs-mediated STAT1 phosphorylation (Figure 7H), while retaining IFNγ signaling inhibition, thus mirroring IVIg EV functions (Figure 7I). These findings highlight the dual actions of IFNγ-associated EVs capable of both activating and inhibiting JAK/STAT1 signaling. Critically, IFNγ-associated IVIg EVs lose the ability to activate IFNγ signaling while retaining their capacity to inhibit it, thereby contributing to the anti-inflammatory effects of IVIg.

We propose that cEP alters IFNγ-bound IVIg EVs, rendering them unable to activate but capable, as decoys, of inhibiting inflammatory pathways (Figure 8). Acute IFNγ triggers receptor dimerization and JAK/STAT1 signaling, whereas chronic exposure downregulates IFNGR1/2 and inhibits signaling (Crisler et al., 2019; Park-Min et al., 2007). cEP likely modifies IFNγ conformation, preventing receptor dimerization and JAK/STAT1 activation, as evidenced by abrogation of STAT1 phosphorylation by cEP IFNγ-eEVs. Given IFNγ’s role in macrophage polarization and adaptive inflammation, this attenuates excessive inflammation, contributing to IVIg’s immunosuppressive effects.

**FIGURE 8.**
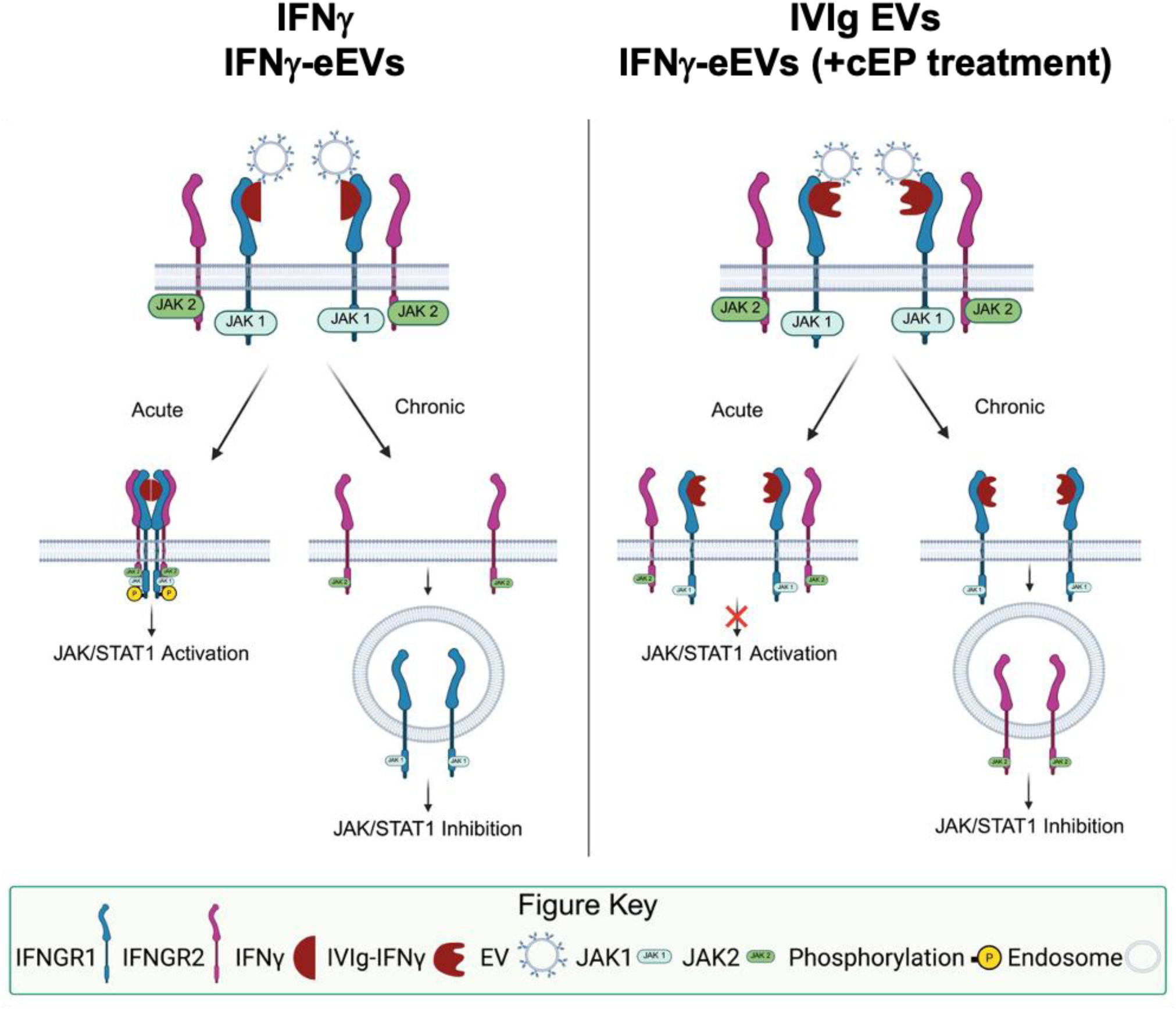
Graphical presentation of IFNγ receptor activation and JAK/STAT1 signaling inhibition by IFNγ-associated EVs. IFNγ is present in both monomers and dimer configuration in solution. IFNγ binding to the IFNGR1/IFNGR2 complex activates JAK/STAT1 signaling (left panel). Similarly, IFNγ binds to the surface of EVs, in part through anti-IFNγ antibodies. The IFNγ-EV complex inhibits IFNGR-mediated JAK/STAT1 activation (right panel). EV, extracellular vesicles; IFNγ, interferon gamma; IFNGR; interferon gamma receptor; JAK, Janus kinase; STAT1, signal transducer and activator of transcription 1.

This study is the first to demonstrate EVs in IVIg preparations and that IFNγ-carrying IVIg EVs strongly suppress IFNGR/JAK/STAT signaling. These anti-inflammatory properties, at least for IFNγ-associated EVs, tied to cEP processing, reveal a novel mechanism for functional modification of EVs in therapeutic plasma products.

## 4 Discussion

The IVIg manufacturing process involves sequential precipitation, fractionation, and purification steps designed to isolate concentrated IgG from pooled human plasma while ensuring pathogen safety and product consistency (Burnouf and Radosevich, 2003; Troccoli et al., 1998). Key stages include cEP, polyclonal IgG enrichment using immunoaffinity chromatography or octanoic acid fractionation coupled with anion exchange chromatography, and nanofiltration (15-35 nm cutoff), to remove viruses and contaminants.

Despite these stringent processes, substantial evidence now demonstrates that intact EVs, including CD63+ populations exceeding the nominal nanofiltration cutoff, persist in multiple IVIg and SCIg preparations. EVs isolated using dUC or SEC, were validated as structurally intact, biologically competent particles by NTA, electron microscopy, and two independent flow cytometry platforms. These findings support the robustness and reproducibility of EV detection in highly processed biologics.

These observations challenge the prevailing assumption that large vesicular structures are effectively eliminated during biopharmaceutical processing. Instead, they suggest that EVs represent a process-resistant component of plasma-derived therapeutics, with potential functional relevance across inflammatory and autoimmune indications.

cEP disrupts plasma membrane architecture and permeability (Bangham et al., 1965; Gutknecht and Tosteson, 1970; Hunt and Kaszuba, 1989; Ly and Longo, 2004), and our data indicate that such disruption contributes to altered EV biophysical and molecular properties. IVIg- and SCIg-derived EVs display increased size distributions, modified membrane integrity, and distinct protein compositions relative to EVs from UHP. Multiplex phenotyping revealed significantly reduced expression of canonical tetraspanins CD9 and CD81, with enrichment of CD63, indicating selective retention or enrichment of a distinct EV subpopulation during manufacturing. This shift in tetraspanin profile is consistent with altered endosomal trafficking and vesicle biogenesis pathways, possibly reflecting stress-induced vesiculation during cEP. Reduced CD9 expression has been linked to CD63 upregulation, and enhanced EV release (Suárez et al., 2017), suggesting coordinated remodeling of vesicle identity.

Importantly, IVIg-derived EVs exhibited diminished expression of HLA-ABC and HLA-DR/DP/DQ markers relative to UHP EVs. This feature may attenuate immunogenicity and contribute to the well-established IVIg anti-inflammatory effects. Concurrently, IVIg EVs were significantly enriched in markers associated with progenitor or “stemness” phenotypes (CD29, ROR1, CD326, CD133/1, and CD24}, likely reflecting selective incorporation during vesicle formation (Geng et al., 2013; Jaggupilli and Elkord, 2012; Joseph et al., 2019; Zhu et al., 2016). While the functional significance of these markers in EV-mediated immune modulation remains to be fully defined, their presence suggests the importance of EV heterogeneity in shaping immunological outcomes. Collectively, these findings support the concept that IVIg processing generates a unique and functionally distinct EV subpopulation rather than preserving native plasma vesicles.

The broad therapeutic efficacy of IVIg in both primary immunodeficiencies and autoimmune diseases is attributed to its capacity to modulate diverse and sometimes opposing immune pathways (Velikova et al., 2023). Our data extend this paradigm by implicating EV-associated cytokine signaling as a contributing mechanism. Multiplex cytokine profiling revealed elevated levels of IFNγ in IVIg relative to UHP. Although classically pro-inflammatory, IFNγ also exerts context-dependent immunoregulatory functions, including suppression of type 2 responses and induction of anti-inflammatory programs (Mühl and Pfeilschifter, 2003). We propose that EV-associated IFNγ is a key mediator of IVIg-induced immune modulation. IVIg-derived EVs demonstrated markedly reduced levels of pro-inflammatory cytokines (e.g., RANTES) compared to UHPs, while anti-inflammatory cytokines such as IL-10 were preferentially detected in non-vesicular protein-rich fractions. This partitioning suggests selective cytokine association with EVs, shaping localized immune signaling environments. Such selective cytokine packaging likely contributes to IVIg efficacy to attenuate pathogenic inflammation across a wide spectrum of disorders.

Cytokine-EV interactions are increasingly recognized as dynamic and affinity-dependent, with many proinflammatory species showing relatively low-affinity binding (Hussain et al., 2020). The reduced pro-inflammatory cytokine levels observed in IVIg EVs are therefore consistent with an overall anti-inflammatory milieu. EVs can function as cytokine reservoirs or decoys, modulating receptor engagement and downstream signaling (Chen et al., 2025; Liang et al., 2023b). In this context, both native IVIg EVs and engineered cEP-derived IFNγ-bearing EVs, could inhibit IFNγ-induced STAT1 activation highlighting a promising generalizable mechanism of immune regulation with translational relevance.

Integrating these findings, we propose a mechanistic model in which IVIg-derived EVs act as natural cytokine decoys that bind and present IFNγ in a controlled vesicle-associated form. This presentation enables sustained, low-level stimulation of myeloid cells, leading to ligand-dependent downregulation of the IFNγ receptor (IFNGR1). Mechanistically, this may involve epigenetic remodeling at regulatory regions of the IFNGR1 locus (Crisler et al., 2019; Cui et al., 2021; Penix et al., 1996; Yu et al., 2025), including reduced H3K4me3 occupancy, and transcriptional repression. As a result, downstream JAK/STAT signaling is attenuated, diminishing STAT1 and STAT3 phosphorylation, ameliorating myeloid cell sensitivity to further IFNγ exposure (Crisler et al., 2019). In autoimmune and inflammatory settings, this pathway may recalibrate excessive IFNγ-driven macrophage activation and suppress pathogenic cytokine cascades.

This model aligns with broader principles of ligand-induced receptor regulation and highlights the unique capacity of EVs to deliver spatially and temporally controlled signals. Unlike soluble cytokines, EV-associated ligands may achieve targeted, prolonged receptor engagement without widespread activation. EV biodistribution may also be influenced by interactions with circulating cells such as erythrocytes and platelets, facilitating targeted delivery and limiting off-target effects (Dai et al., 2022; Livkisa et al., 2024; Pavlova et al., 2026).

From a translational perspective, this framework suggests that these EVs are not merely nonfunctional contaminants of IVIg manufacturing but actively contribute to its clinical efficacy. This raises the possibility that eEVs, generated through controlled processes such as cEP, could be developed as targeted immunomodulatory agents. Such approaches may enable precision therapies for IFNγ-driven disorders, including multiple sclerosis, Guillain-Barré syndrome and other chronic inflammatory disorders (Nimmerjahn and Ravetch, 2007; Sorensen, 2003; van Doorn et al., 2010).

In conclusion, CD63-enriched EVs persist in IVIg preparations despite extensive purification and exhibit a distinct molecular signature characterized by reduced CD9/CD81 and HLA expression, enrichment of stemness-associated markers, and selective association with IFNγ. These EVs function as bioactive immunomodulators capable of sequestering cytokines, modulating receptor signaling, and attenuating inflammatory responses. By linking EV-mediated cytokine regulation to epigenetic control of inflammatory signaling pathways, this work proposes a unifying mechanism underlying the dual immunostimulatory and immunosuppressive effects of IVIg.

Future studies should focus on comprehensive characterization of EV cargo (including proteins, RNAs, and biomolecular corona components, delineation of EV-immune cell interactions in vivo, and validation of these mechanisms in disease models. Such efforts will advance understanding of IVIg and IFNγ biology and support the development of next-generation EV-based therapeutics for inflammatory and autoimmune disorders, potentially reducing reliance on traditional immunoglobulin replacement therapies.

## Author contributions

**Conceptualization:** MGM, APP, AM, LAH, JCJ; **Investigation**: KK, KEP, CRS, AA, AP, SMC, MLP, JG, RDC, AES, AJL, GFD, PJW, AM, JCJ, LAH, APP, MGM; **Funding acquisition:** MGM, APP, LAH, JCJ; **Supervision:** MGM, LAH, JCJ, APP; **Resources:** MGM, APP, LAH, AJL **Writing – original draft:** MGM, APP, PJW; **Writing – review & editing:** KK, KEP, CRS, AA, AP, SMC, MLP, JG, RDC, AES, AJL, GFD, PJW, AM, JCJ, LAH, APP, MGM

## Acknowledgements

The authors would like to thank Dr. R. John Looney and Dayle Ewanow, RN (Dept of Medicine: Allergy, Immunology, and Rheumatology, U of Rochester Medical Center, Rochester, NY) for supplying the IVIg and SCIg.

## Data Availability Statement

No large datasets were generated or analyzed during this study. The data that support the findings of this study are available on request from the corresponding author.

## Abbreviations

BC: Biomolecular corona
BCA: bicinchoninic acid
CD: Cluster of Differentiation
EV: Extracellular vesicle
Fc: Fragment crystallizable
HLA: Human leukocyte antigen
IFNGR1: Interferon gamma receptor 1
IFNGR2: Interferon gamma receptor 2
IFNγ: Interferon gamma
Ig: Immunoglobulin
IgRT: Immunoglobulin replacement therapy
IL: Interleukin
IO: Immuno-oncology
IVIg: Intravenous immunoglobulin
JAK: Janus kinase
NTA: nanoparticle tracking analysis
p-STAT1: Phosphorylated signal transducer and activator of transcription 1
SCIg: Subcutaneous immunoglobulin
SEC: size-exclusion chromatography
STAT1: Signal transducer and activator of transcription 1
TEM: transmission electron microscopy
UHPi: Individual unprocessed human plasma
UHPp: Pooled unprocessed human plasma.

## Conflict of interest

The authors have declared that no conflict of interest exists.

## Funding

This work was supported by grants from the National Institutes of Health to MGM (R01-AR074314; R21-DE033558); and the Kimmel Cancer Center Support Grant (5P30CA056036-20) to the Thomas Jefferson University Flow Cytometry and Human Immune Monitoring Core.

## Supplemental Figures

**FIGURE S1.** TEM images of IVIg EVs. (A) High magnification of TEM images of IVIg EVs (size bar = 100 nm). (B) Representative low magnification of TEM images of multiple IVIg EVs (size bar = 600 nm). (C) Composite of fifteen TEM images of IVIg EVs ranging in size from approximately 50 nm to >200 mm in diameters. EV, extracellular vesicles; IVIg, intravenous immunoglobulin; TEM, transmission electron microscopy.

**FIGURE S2.** CD63 positive IVIg and SCIg EVs. Immunoblotting analyses of UHPi, IVIg, and SCIg void (1-4), EV-rich (5-9), and protein-rich (10-14) fractions for expression of the small EV-enriched marker CD63 and human IgG. EV, extracellular vesicles; IgG, immunoglobulin G; IVIg, intravenous immunoglobulin; SCIg, subcutaneous immunoglobulin; SEC, size-exclusion chromatography; UHPi, individual unprocessed human plasma.

**FIGURE S3.** Detection of CD63^+^ EVs in IVIg by flow cytometry. EVs were labeled with DiD or CD63-PE. (A) Gating strategy for fluorescent EVs by conventional flow cytometry. Scatter plots (log scale) of unlabeled (left) or DiD-labeled EVs (right). (B) Histogram plots of CD63 expression on DiD^+^ events (gated on R1). Isotype control (blue) indicates background; representative CD63 staining shown for UHPp EVs (orange) and IVIg EVs (red). (C) Representative imaging flow cytometry images of EVs showing BF, CD63, DiD, and scatter channels. BF, bright field; DiD, 1,1′-dioctadecyl-3,3,3′,3′-tetramethylindodicarbocyanine, 4-chlorobenzenesulfonate salt; EV, extracellular vesicles; IVIg, intravenous immunoglobulin; UHPp, pooled unprocessed human plasma.

**FIGURE S4.** Comparison of surface‒associated cytokines in UHPi, IVIg, and SCIg SEC fractions. UHPi, IVIg and SCIg were purified over SEC and fifteen fractions were collected. Representative fractions 4-14 were plotted as bar graphs showing cytokines, chemokines, and growth factors that were associated with the EV-rich (A), EV-rich and protein-rich (B), protein-rich (C), and not detectable (D). EV, extracellular vesicles; Ig, immunoglobulin; IVIg, intravenous immunoglobulin; SCIg, subcutaneous immunoglobulin; SEC, size-exclusion chromatography; UHPi, individual unprocessed human plasma.

**FIGURE S5.** Lower inflammatory nature of IVIg EV-rich fractions. EV-rich SEC fractions 5-9 from different lots of UHPi (n=6) and IVIg (n=3) were subjected to Luminex analysis for cytokines, chemokines, and growth factors. Twenty-two cytokines were present at significantly lower levels (A), while eight were present at higher levels (B), in IVIg compared with UHPi. EVs, extracellular vesicles; IVIg, intravenous immunoglobulin; SEC, size-exclusion chromatography; UHPi, individual unprocessed human plasma. P-value: * P ≤ 0.05.

**FIGURE S6.** Cytokine profiling of EVs derived from UHPi compared to UHPp. (A-B) EVs were purified from 3 individual preps of UHPi and 3 pooled preps of UHPp. The EV-rich fractions (F5-9) were subjected to Luminex analysis for the levels of cytokines, chemokines, and growth factors. Cytokines showing no difference (A) or statistically significant difference (B) but under the limit of detection (C). P-value: ns P > 0.05; * P ≤ 0.05; ** P ≤ 0.01. EV, extracellular vesicles; UHPi, individual unprocessed human plasma; UHPp, pooled unprocessed human plasma.

**FIGURE S7.** Cytokine profiling of EVs derived from SCIg compared to UHPi. EVs were purified from SCIg and UHPi. The EV-rich fractions (F5-9) were subjected to Luminex analysis for the levels of cytokines, chemokines, and growth factors. (A) With the exception of TNFβ, a similar decrease in cytokines was observed in SCIg. (B) Also similar to IVIg, IFNγ was higher in SCIg compared to UHPi. EV, extracellular vesicles; IVIg, intravenous immunoglobulin; SCIg, subcutaneous immunoglobulin; TNFβ, tumor necrosis factor-beta; UHPi, individual unprocessed human plasma.

**FIGURE S8.** Cytokine profiling of the protein-rich fractions of IVIg compared to UHPi. IVIg (n=3) and UHPi (n=6) were purified as described above. Protein-rich fractions (F10-14) of IVIg and UHPi were combined, concentrated, and subjected to Luminex analysis for cytokine level quantification. Twelve cytokines displayed lower levels (A) and seven displayed higher levels (B) in IVIg compared to UHPi. IVIg, intravenous immunoglobulin; UHPi, individual unprocessed human plasma; P-value: * P ≤ 0.05.

**FIGURE S9.** IFNγ binds to the surface of EVs, in part, through anti-IFNγ antibodies. Western blotting analysis of recombinant IFNγ (1 and 0.1 µg/lane) alone or treated with β-mercaptoethanol for anti-IFNγ antibody using unfractionated (top panels), EV-rich (middle panels), and protein-rich (bottom panels) fractions of UHPp, UHPi, IVIg, and SCIg. Lane 1=1 µg/lane; Lane 2=0.1 µg/lane. β-mer, β-mercaptoethanol; EVs, extracellular vesicles; IFNγ, interferon gamma; IVIg, intravenous immunoglobulin; SCIg, subcutaneous immunoglobulin; UHPi, individual unprocessed human plasma; UHPp, pooled unprocessed human plasma.

**FIGURE S10.** UHPi and UHPp did not inhibit IFNγ-mediated p-STAT1 activation. EVs were purified from 10 mL each of UHPi (solid circle) and IVIg (open circle) by SEC using Izon qEV10 columns. (A) Seven 5 mL fractions were collected, and protein concentrations were determined by BCA assay. (B) Immunoblotting for RANTES and IgG. (C) Cells were incubated with high doses (10 - 30 µg/mL) of UHPp and UHPi EVs for 24 h in serum free medium and then were treated with recombinant IFNγ (3 ng/mL) for 15 min. Cells were lysed and proteins were immunoblotted for p-STAT1, STAT1, β-Actin. (D) Cells were incubated with high doses (10 - 30 µg/mL) of UHPp and UHPi EVs for 24 h in serum free medium and then were treated with recombinant IFNγ (3 ng/mL) for 15 min. Cells were lysed and proteins were immunoblotted for p-STAT1, STAT1, and β-Actin. BCA, bicinchoninic acid assay; EV, extracellular vesicles; IFNγ, interferon gamma; Ig, immunoglobulin; IVIg, intravenous immunoglobulin; RANTES, regulated upon activation normal T cell expressed and secreted; p-STAT1, phosphorylated signal transducer and activator of transcription 1; p-STAT3, signal transducer and activator of transcription 3; STAT1, signal transducer and activator of transcription 1; STAT3, signal transducer and activator of transcription 3; UHPi, individual unprocessed human plasma; and UHPp, pooled unprocessed human plasma.

**FIGURE S11.** IVIg EV-mediated functions retained after lipoprotein depletion. (A). Immunoblotting of UHPp and IVIg EVs for Apo A1, Apo B, and IgG. (B) For activation, cells were stimulated with recombinant IFNγ (3 ng/mL), IVIg EVs (9 µg/mL), and LipoMin-treated IVIg EVs (9 µg/mL) for 15 min and proteins lysed and immunoblotted for p-STAT1, STAT1, and β-Actin. (C) For inhibition, cells were pre-incubated with IVIg EVs or LipoMin-treated IVIg EVs for 24 h, then stimulated with recombinant IFNγ (3 ng/mL) for 15 min, and immunoblotted for p-STAT1, STAT1, and β-Actin. EV, extracellular vesicles; IVIg, intravenous immunoglobulin; IVIg, intravenous immunoglobulin; IFNγ, interferon gamma; p-STAT1, phosphorylated signal transducer and activator of transcription 1; STAT1, signal transducer and activator of transcription 1.

## Notes

### Competing Interest Statement

The authors have declared no competing interest.

