## Supplemental Figures for "Translational Opportunity of Engineered IFNγ-eEVs Through Targeted Inhibition of JAK/STAT1 Signaling, Mimicking IVIg Therapy"

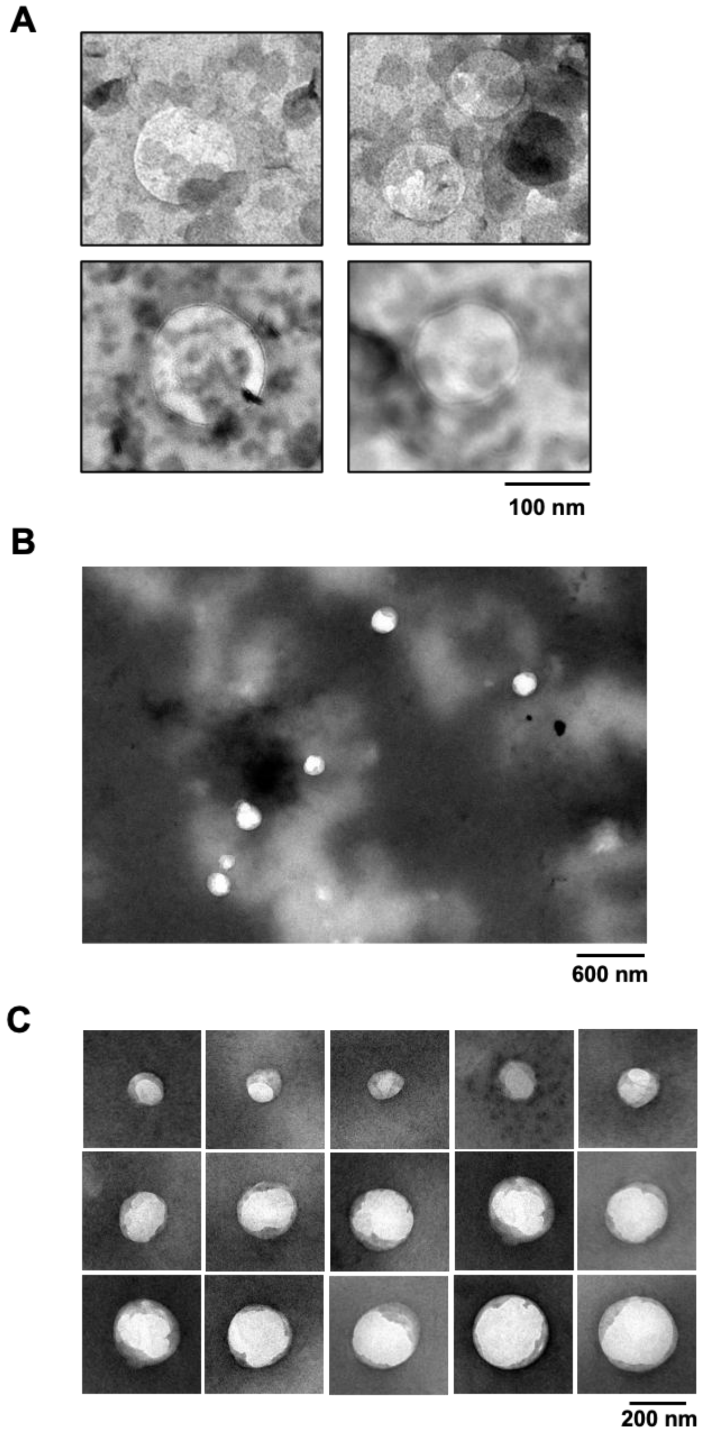


**FIGURE S1.** TEM images of IVIg EVs. (A) High magnification of TEM images of IVIg EVs (size bar = 100 nm). (B) Representative low magnification of TEM images of multiple IVIg EVs (size bar = 600 nm). (C) Composite of fifteen TEM images of IVIg EVs ranging in size from approximately 50 nm to >200 mm in diameters. EV, extracellular vesicles; IVIg, intravenous immunoglobulin; TEM, transmission electron microscopy.

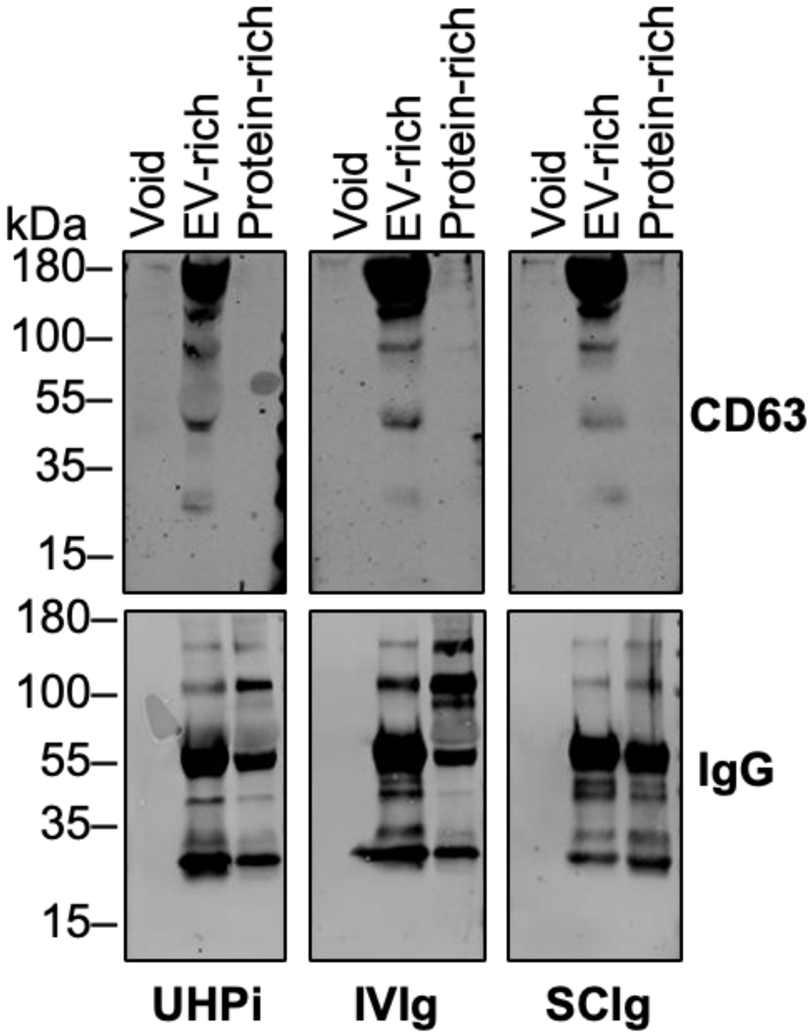


**FIGURE S2.** CD63 positive IVIg and SCIg EVs. Immunoblotting analyses of UHPi, IVIg, and SCIg void (1-4), EV-rich (5-9), and protein-rich (10-14) fractions for expression of the small EV-enriched marker CD63 and human IgG. EV, extracellular vesicles; IgG, immunoglobulin G; IVIg, intravenous immunoglobulin; SCIg, subcutaneous immunoglobulin; SEC, size-exclusion chromatography; UHPi, individual unprocessed human plasma.

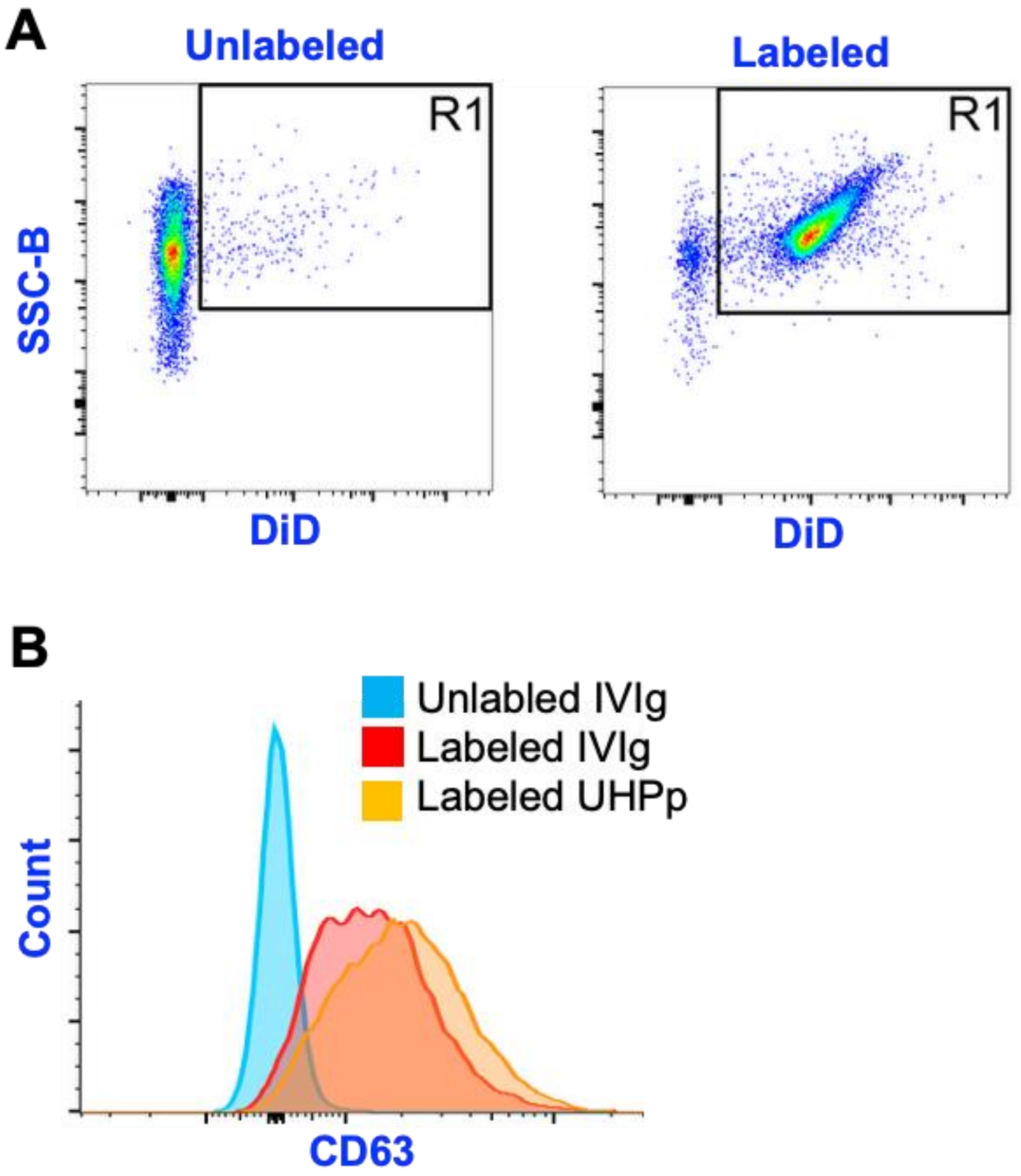


**FIGURE S3.** Detection of CD63$^{+}$ EVs in IVIg by flow cytometry. EVs were labeled with DiD or CD63-PE. (A) Gating strategy for fluorescent EVs by conventional flow cytometry. Scatter plots (log scale) of unlabeled (left) or DiD-labeled EVs (right). (B) Histogram plots of CD63 expression on DiD$^{+}$ events (gated on R1). Isotype control (blue) indicates background; representative CD63 staining shown for UHPp EVs (orange) and IVIg EVs (red). (C) Representative imaging flow cytometry images of EVs showing BF, CD63, DiD, and scatter channels. BF, bright field; DiD, 1,1′-dioctadecyl-3,3,3′,3′-tetramethylindodicarbocyanine, 4-chlorobenzenesulfonate salt; EV, extracellular vesicles; IVIg, intravenous immunoglobulin; UHPp, pooled unprocessed human plasma.

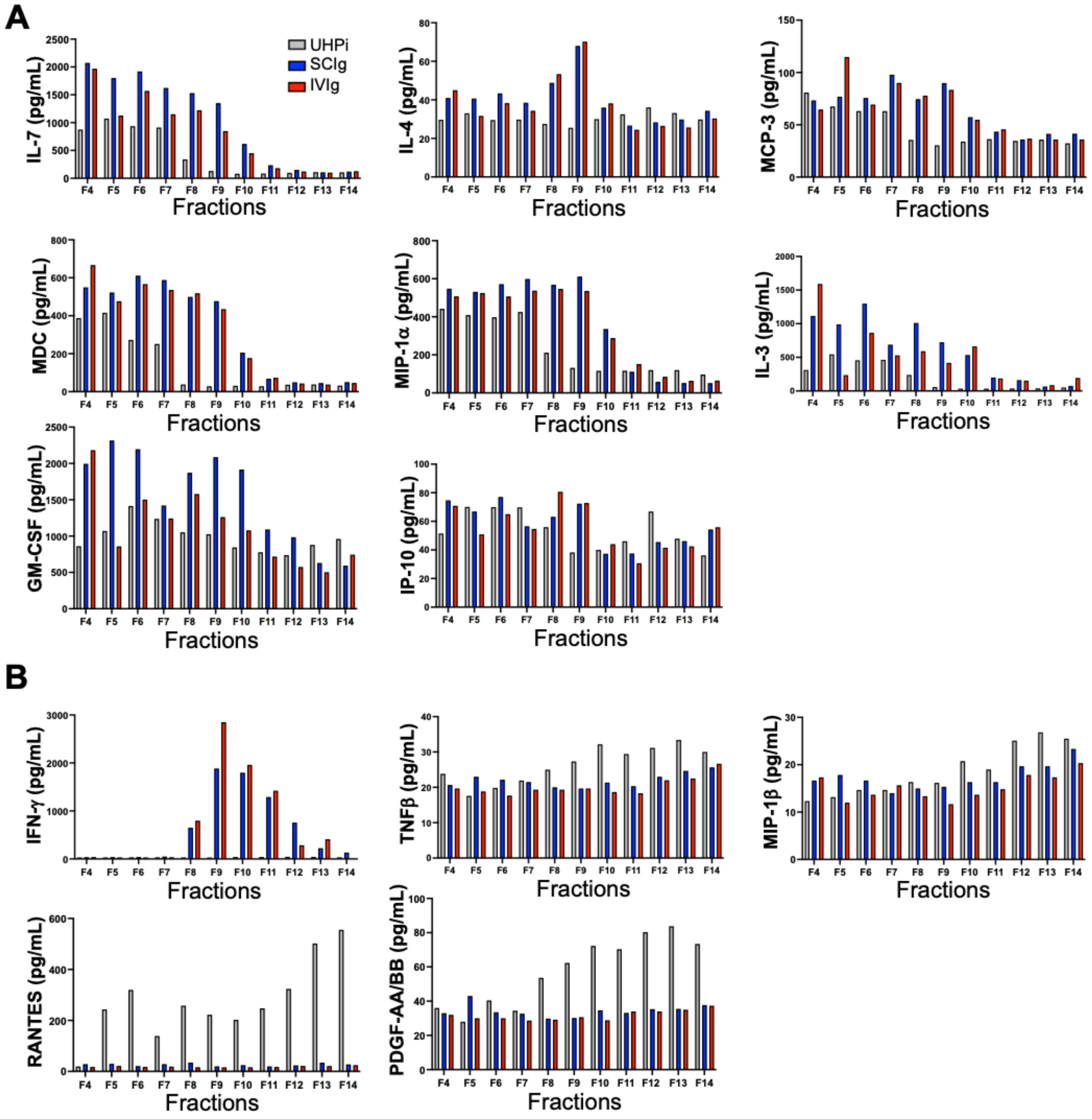


**FIGURE S4.** Comparison of surface–associated cytokines in UHPi, IVIg, and SCIg SEC fractions. UHPi, IVIg and SCIg were purified over SEC and fifteen fractions were collected. Representative fractions 4-14 were plotted as bar graphs showing cytokines, chemokines, and growth factors that were associated with the EV-rich (A), EV-rich and protein-rich (B), protein-rich (C), and not detectable (D). EV, extracellular vesicles; Ig, immunoglobulin; IVIg, intravenous immunoglobulin; SCIg, subcutaneous immunoglobulin; SEC, size-exclusion chromatography; UHPi, individual unprocessed human plasma.

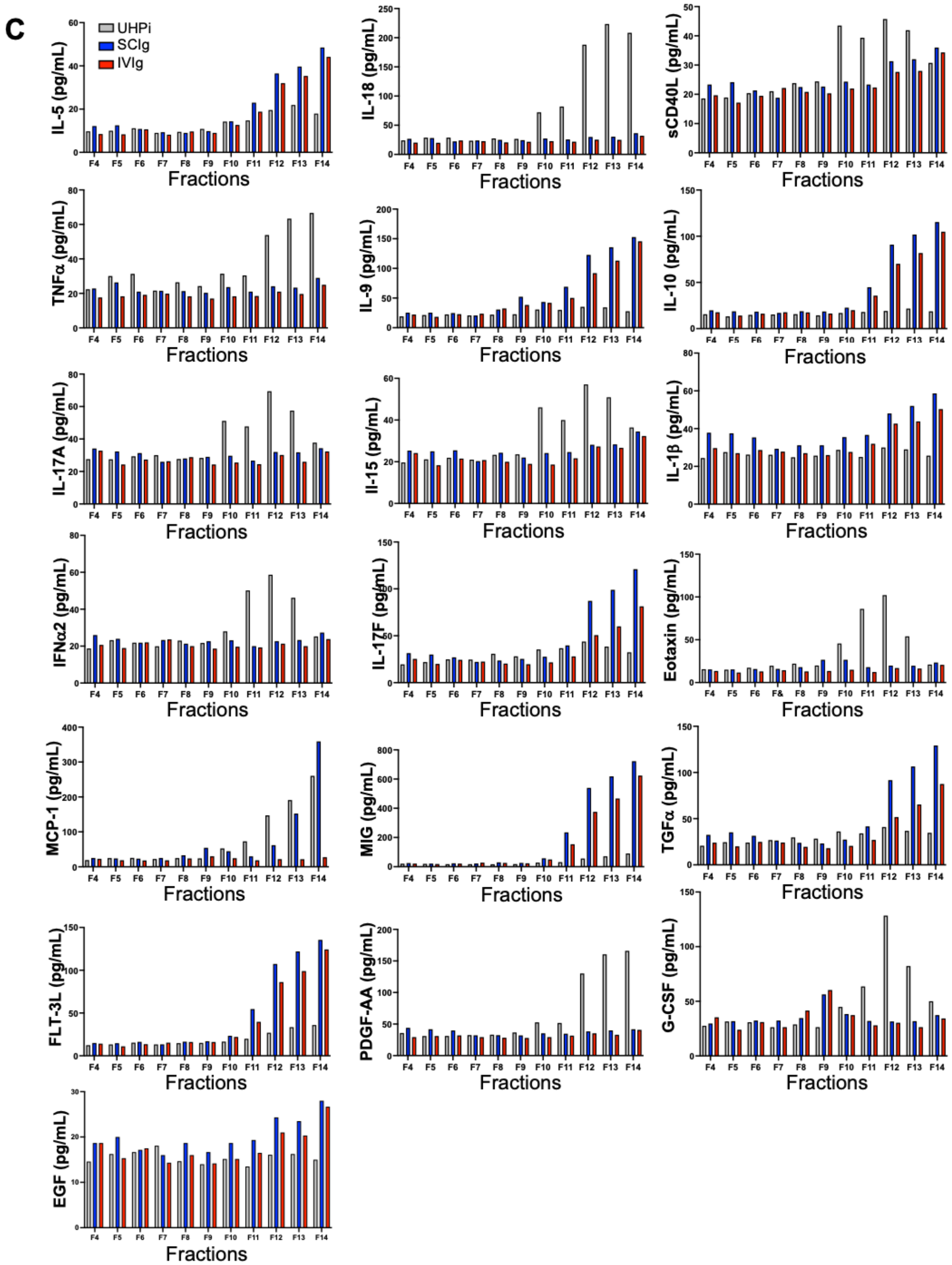


**FIGURE S4.** Comparison of surface–associated cytokines in UHPi, IVIg, and SCIg SEC fractions. UHPi, IVIg and SCIg were purified over SEC and fifteen fractions were collected. Representative fractions 4-14 were plotted as bar graphs showing cytokines, chemokines, and growth factors that were associated with the EV-rich (A), EV-rich and protein-rich (B), protein-rich (C), and not detectable (D). EV, extracellular vesicles; Ig, immunoglobulin; IVIg, intravenous immunoglobulin; SCIg, subcutaneous immunoglobulin; SEC, size-exclusion chromatography; UHPi, individual unprocessed human plasma.

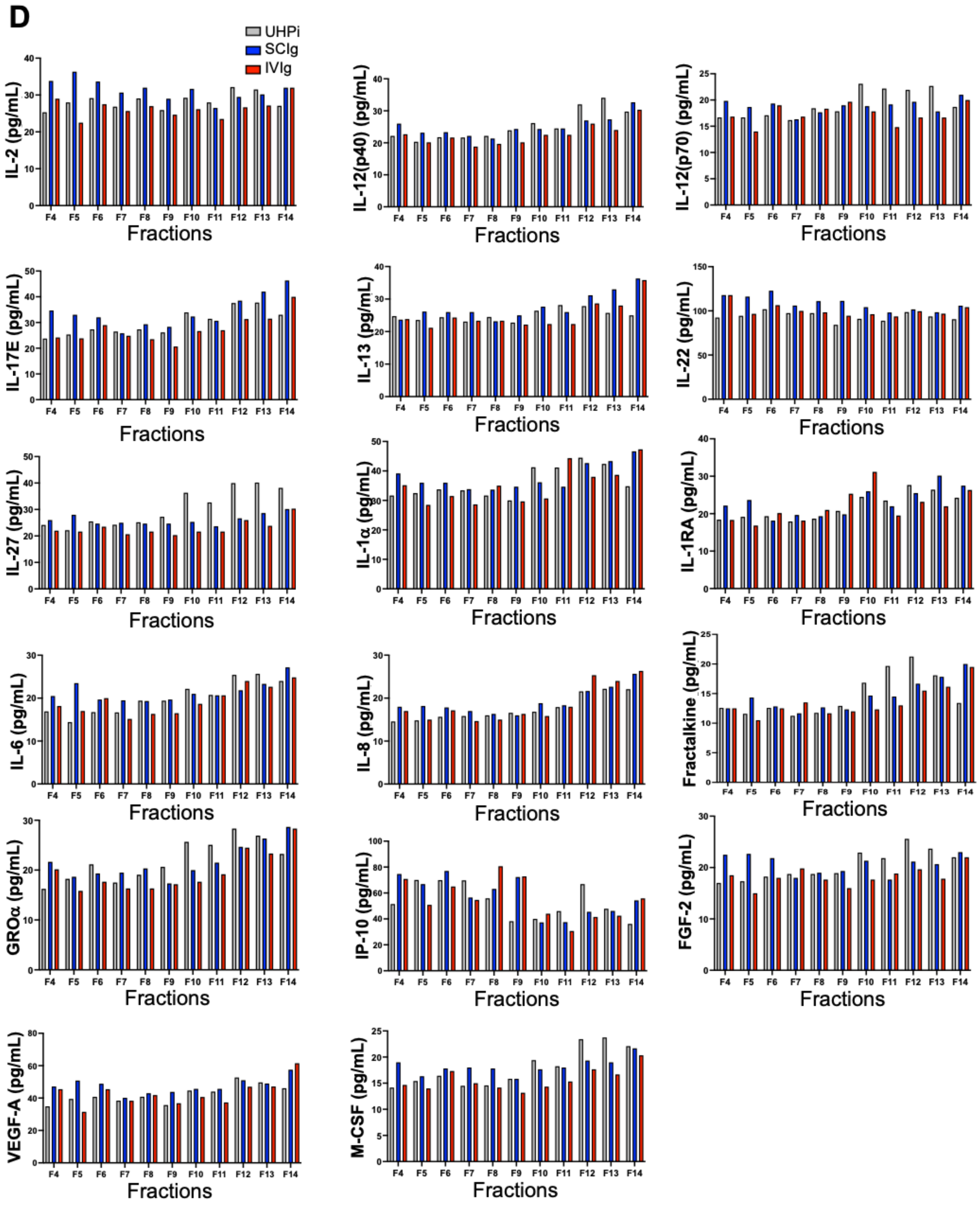


**FIGURE S4.** Comparison of surface–associated cytokines in UHPi, IVIg, and SCIg SEC fractions. UHPi, IVIg and SCIg were purified over SEC and fifteen fractions were collected. Representative fractions 4-14 were plotted as bar graphs showing cytokines, chemokines, and growth factors that were associated with the EV-rich (A), EV-rich and protein-rich (B), protein-rich (C), and not detectable (D). EV, extracellular vesicles; Ig, immunoglobulin; IVIg, intravenous immunoglobulin; SCIg, subcutaneous immunoglobulin; SEC, size-exclusion chromatography; UHPi, individual unprocessed human plasma.

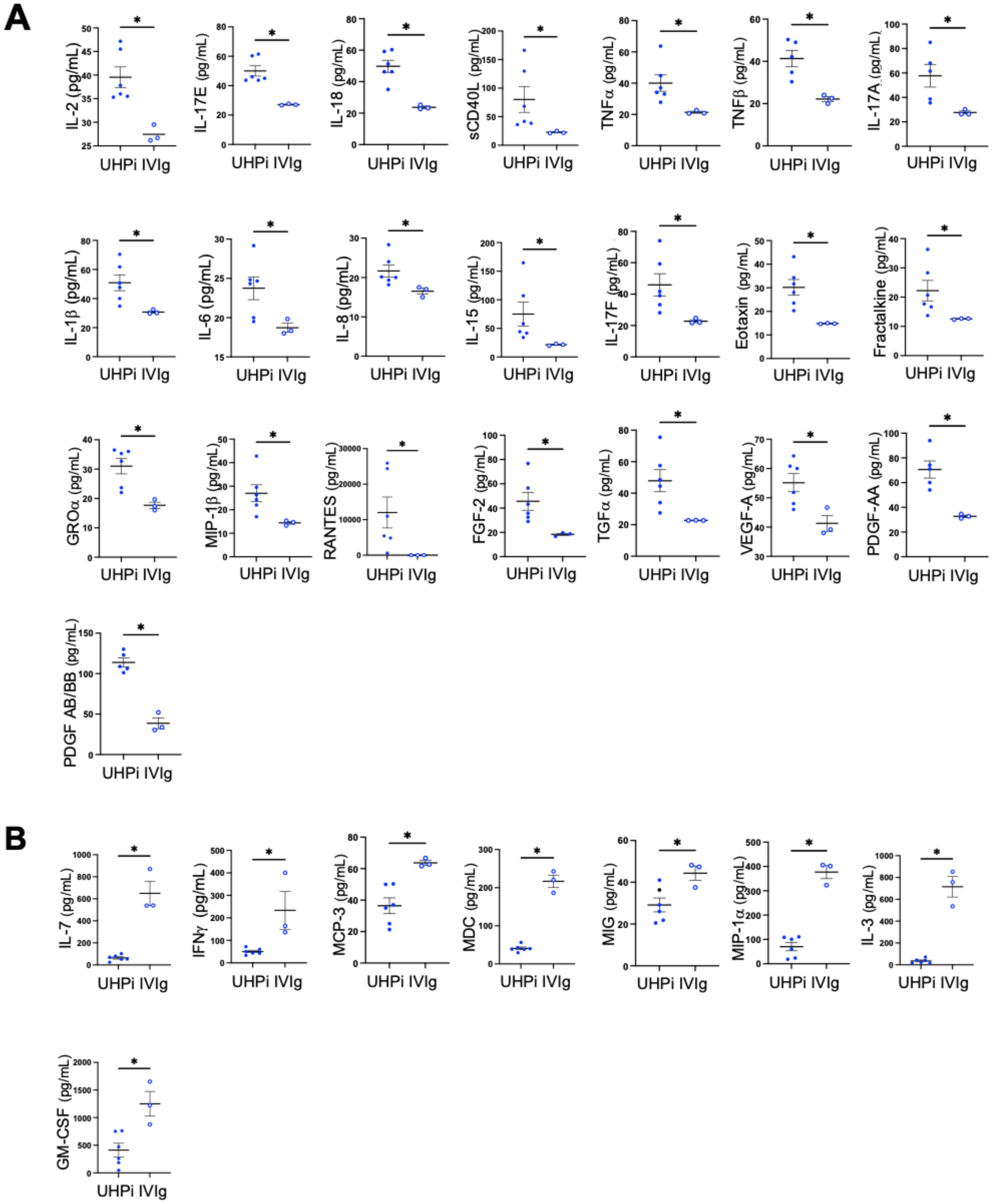


**FIGURE S5.** Lower inflammatory nature of IVIg EV-rich fractions. EV-rich SEC fractions 5-9 from different lots of UHPi (n=6) and IVIg (n=3) were subjected to Luminex analysis for cytokines, chemokines, and growth factors. Twenty-two cytokines were present at significantly lower levels (A), while eight were present at higher levels (B), in IVIg compared with UHPi. EVs, extracellular vesicles; IVIg, intravenous immunoglobulin; SEC, size-exclusion chromatography; UHPi, individual unprocessed human plasma. P-value: * P ≤ 0.05.

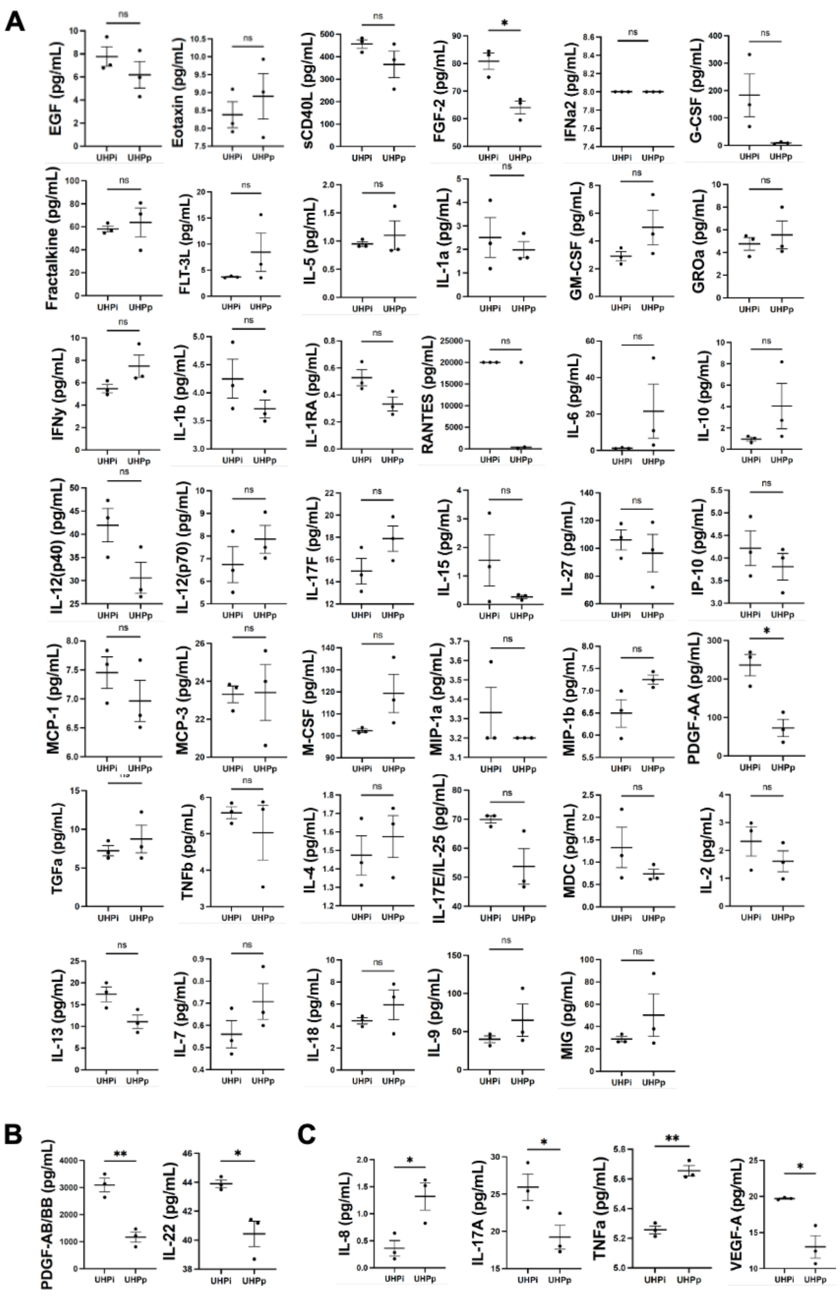


**FIGURE S6.** Cytokine profiling of EVs derived from UHPi compared to UHPp. (A-B) EVs were purified from 3 individual preps of UHPi and 3 pooled preps of UHPp. The EV-rich fractions (F5-9) were subjected to Luminex analysis for the levels of cytokines, chemokines, and growth factors. Cytokines showing no difference (A) or statistically significant difference (B) but under the limit of detection (C). P-value: ns $P > 0.05$; * $P \leq 0.05$; ** $P \leq 0.01$. EV, extracellular vesicles; UHPi, individual unprocessed human plasma; UHPp, pooled unprocessed human plasma.

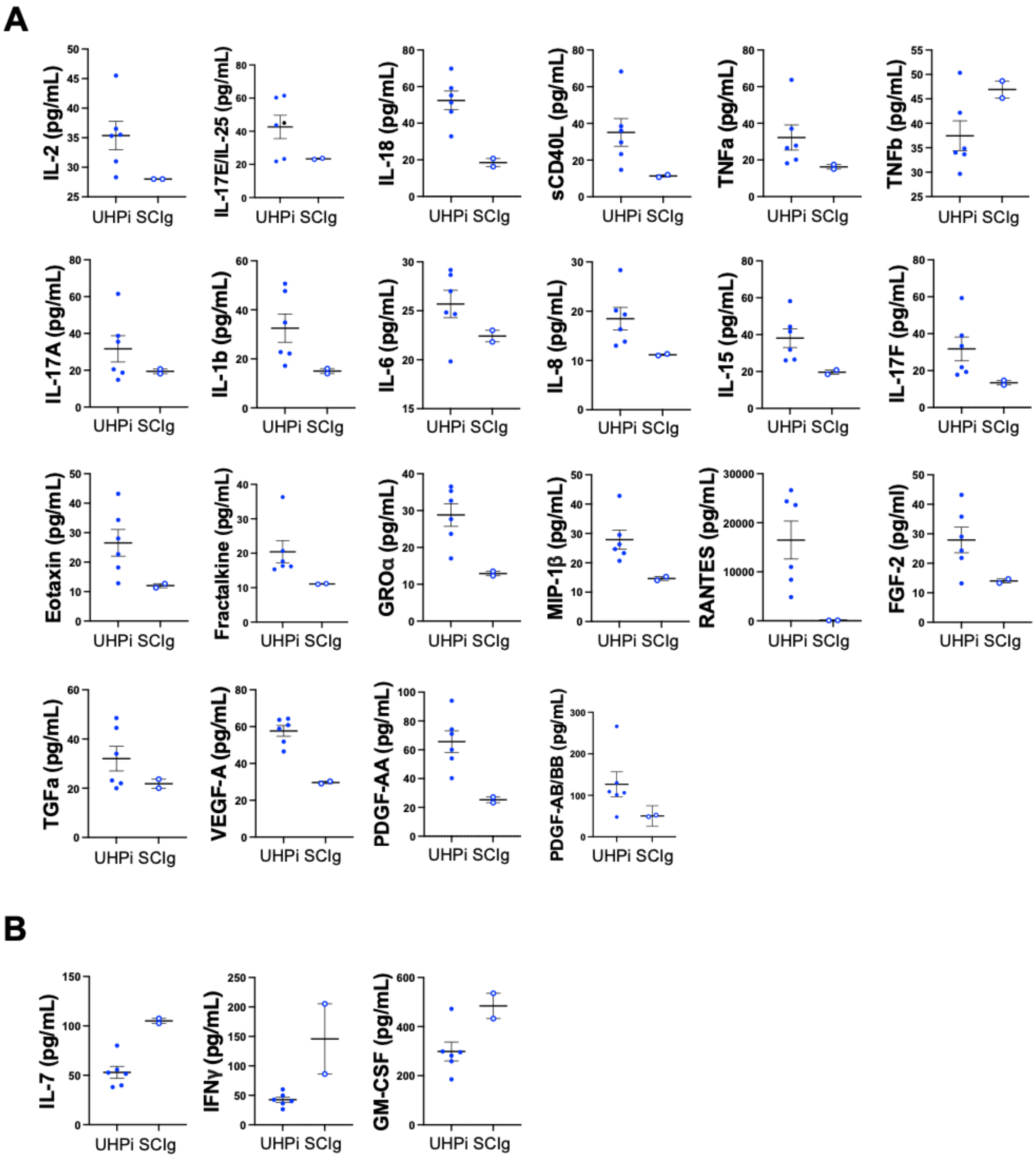


**FIGURE S7.** Cytokine profiling of EVs derived from SCIg compared to UHPi. EVs were purified from SCIg and UHPi. The EV-rich fractions (F5-9) were subjected to Luminex analysis for the levels of cytokines, chemokines, and growth factors. (A) With the exception of TNFβ, a similar decrease in cytokines was observed in SCIg. (B) Also similar to IVIg, IFNγ was higher in SCIg compared to UHPi. EV, extracellular vesicles; IVIg, intravenous immunoglobulin; SCIg, subcutaneous immunoglobulin; TNFβ, tumor necrosis factor-beta; UHPi, individual unprocessed human plasma.

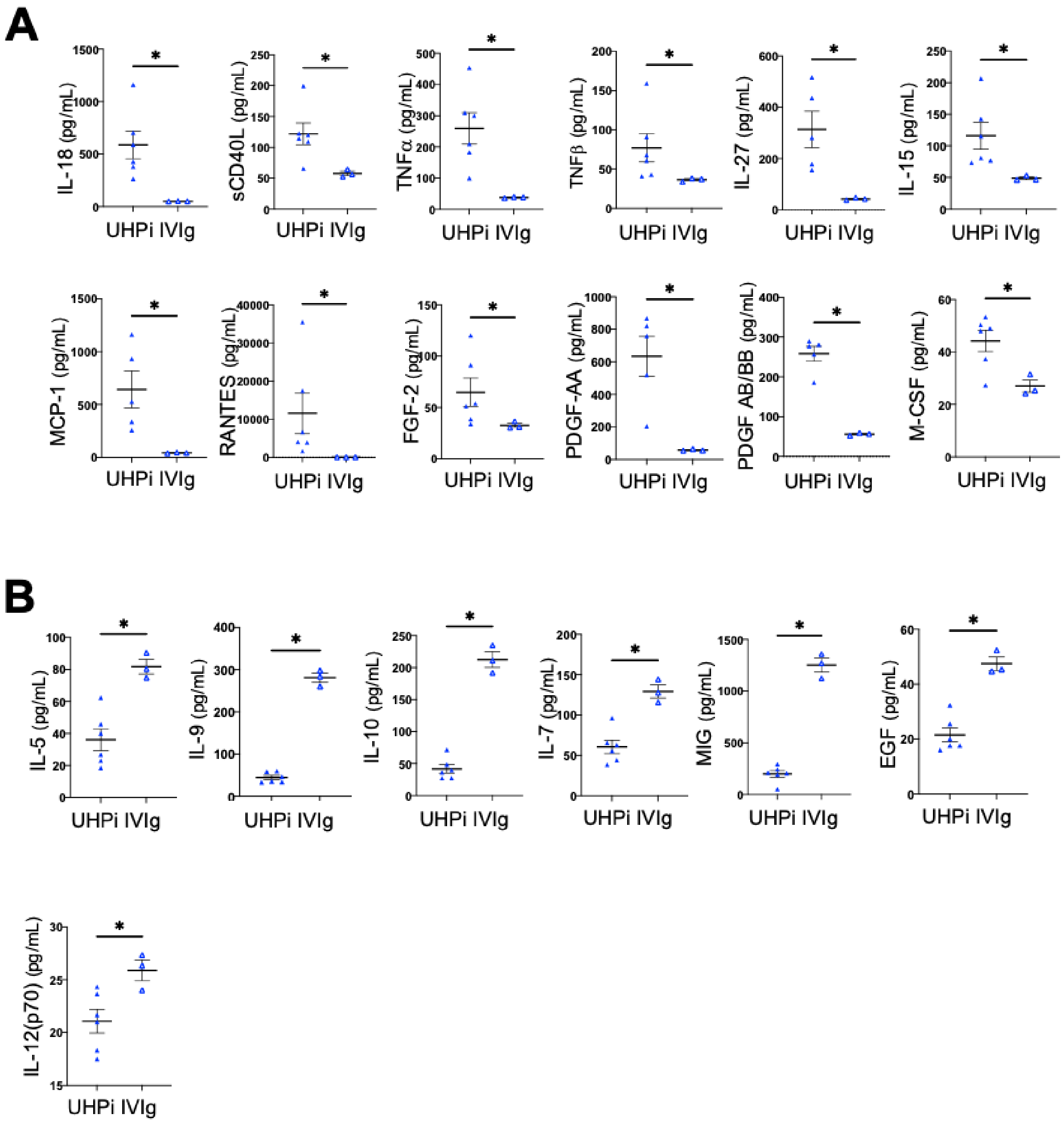


**FIGURE S8.** Cytokine profiling of the protein-rich fractions of IVIg compared to UHPi. IVIg (n=3) and UHPi (n=6) were purified as described above. Protein-rich fractions (F10-14) of IVIg and UHPi were combined, concentrated, and subjected to Luminex analysis for cytokine level quantification. Twelve cytokines displayed lower levels (A) and seven displayed higher levels (B) in IVIg compared to UHPi. IVIg, intravenous immunoglobulin; UHPi, individual unprocessed human plasma; P-value: * P ≤ 0.05.

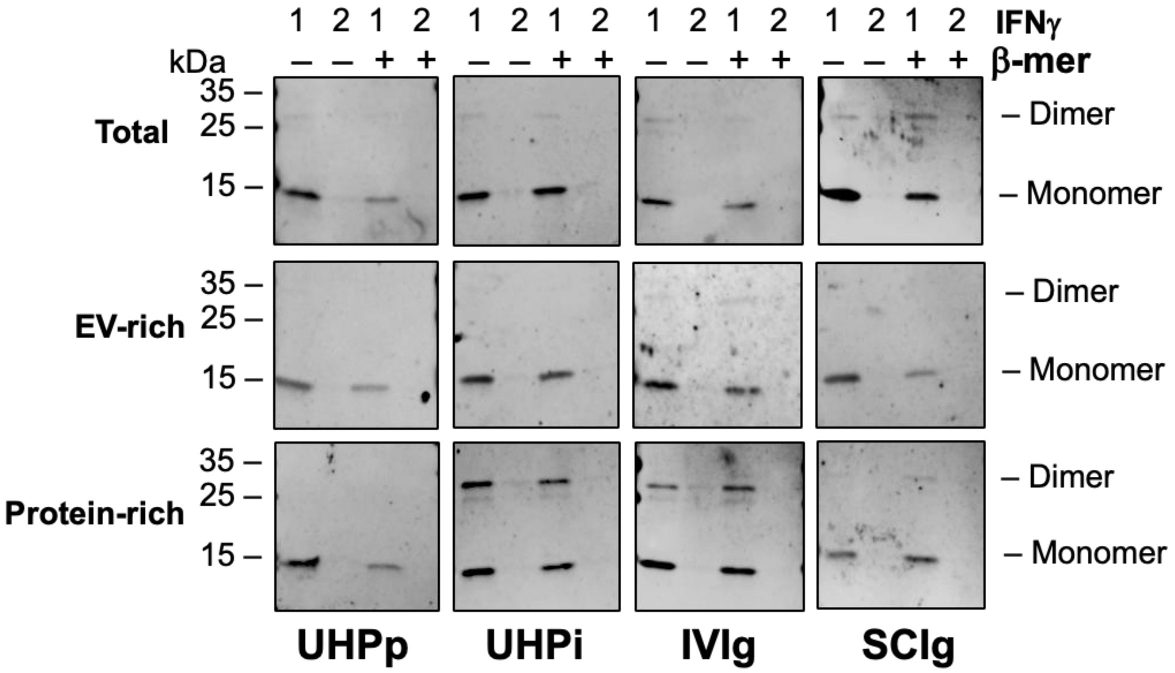


**FIGURE S9.** IFNγ binds to the surface of EVs, in part, through anti-IFNγ antibodies. Western blotting analysis of recombinant IFNγ (1 and 0.1 µg/lane) alone or treated with β-mercaptoethanol for anti-IFNγ antibody using unfractionated (top panels), EV-rich (middle panels), and protein-rich (bottom panels) fractions of UHPp, UHPi, IVIg, and SCIg. Lane 1=1 µg/lane; Lane 2=0.1 µg/lane. β-mer, β-mercaptoethanol; EVs, extracellular vesicles; IFNγ, interferon gamma; IVIg, intravenous immunoglobulin; SCIg, subcutaneous immunoglobulin; UHPi, individual unprocessed human plasma; UHPp, pooled unprocessed human plasma.

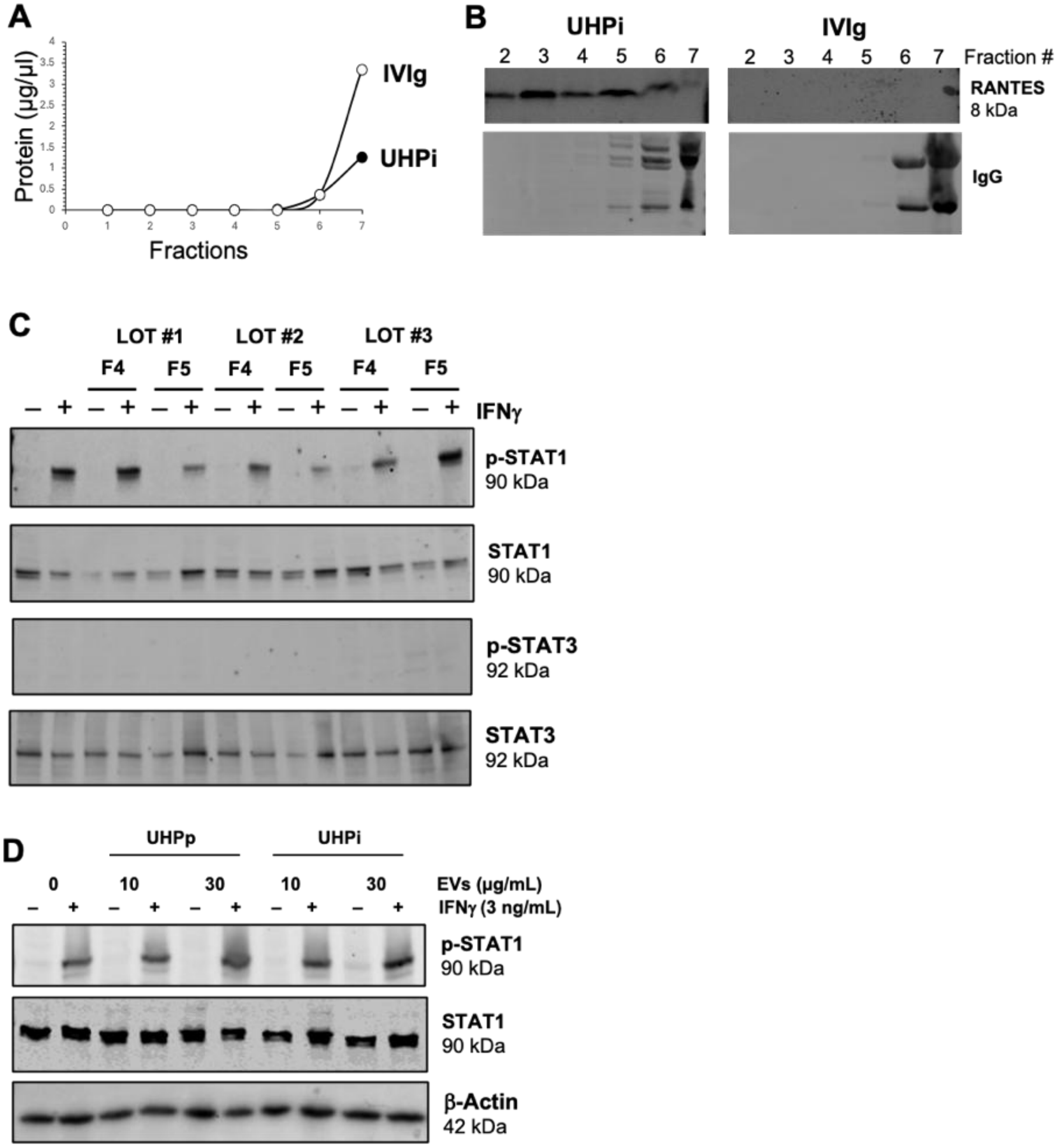


**FIGURE S10.** UHPi and UHPp did not inhibit IFNγ-mediated p-STAT1 activation. EVs were purified from 10 mL each of UHPi (solid circle) and IVIg (open circle) by SEC using Izon qEV10 columns. (A) Seven 5 mL fractions were collected, and protein concentrations were determined by BCA assay. (B) Immunoblotting for RANTES and IgG. (C) Cells were incubated with high doses (10 - 30 µg/mL) of UHPp and UHPi EVs for 24 h in serum free medium and then were treated with recombinant IFNγ (3 ng/mL) for 15 min. Cells were lysed and proteins were immunoblotted for p-STAT1, STAT1, β-Actin. (D) Cells were incubated with high doses (10 - 30 µg/mL) of UHPp and UHPi EVs for 24 h in serum free medium and then were treated with recombinant IFNγ (3 ng/mL) for 15 min. Cells were lysed and proteins were immunoblotted for p-STAT1, STAT1, and β-Actin. BCA, bicinchoninic acid assay; EV, extracellular vesicles; IFNγ, interferon gamma; Ig, immunoglobulin; IVIg, intravenous immunoglobulin; RANTES, regulated upon activation normal T cell expressed and secreted; p-STAT1, phosphorylated signal transducer and activator of transcription 1; p-STAT3, signal transducer and activator of transcription 3; STAT1, signal transducer and activator of transcription 1; STAT3, signal transducer and activator of transcription 3; UHPi, individual unprocessed human plasma; and UHPp, pooled unprocessed human plasma.

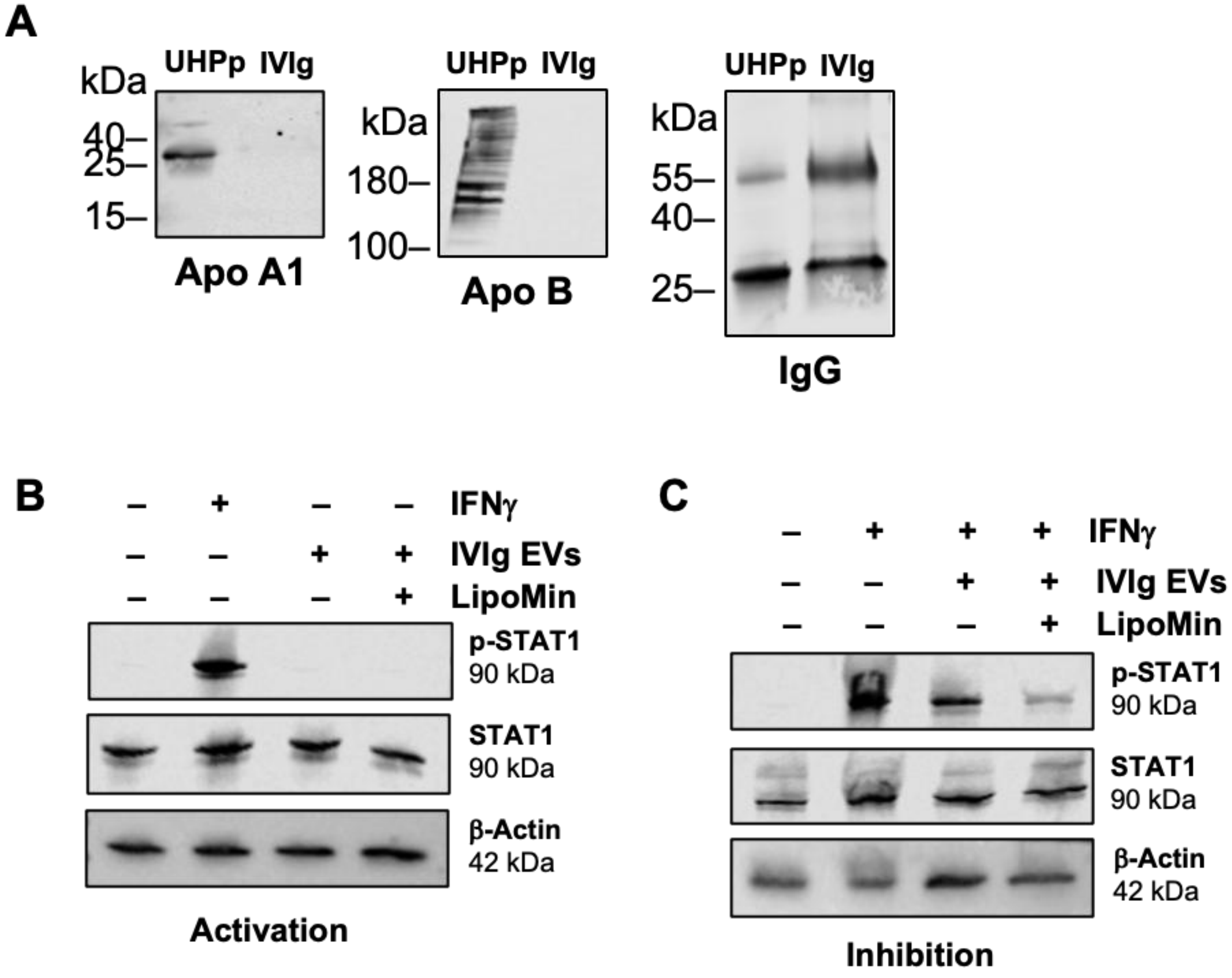


**FIGURE S11.** IVIg EV-mediated functions retained after lipoprotein depletion. (A). Immunoblotting of UHPp and IVIg EVs for Apo A1, Apo B, and IgG. (B) For activation, cells were stimulated with recombinant IFNγ (3 ng/mL), IVIg EVs (9 μg/mL), and LipoMin-treated IVIg EVs (9 μg/mL) for 15 min and proteins lysed and immunoblotted for p-STAT1, STAT1, and β-Actin. (C) For inhibition, cells were pre-incubated with IVIg EVs or LipoMin-treated IVIg EVs for 24 h, then stimulated with recombinant IFNγ (3 ng/mL) for 15 min, and immunoblotted for p-STAT1, STAT1, and β-Actin. EV, extracellular vesicles; IVIg, intravenous immunoglobulin; IVIg, intravenous immunoglobulin; IFNγ, interferon gamma; p-STAT1, phosphorylated signal transducer and activator of transcription 1; STAT1, signal transducer and activator of transcription 1.
